# A computational framework linking molecular regulation, synaptic plasticity, and brain disorders

**DOI:** 10.1101/2025.04.07.647510

**Authors:** Ze-Zheng Li, Wen-Hao Wu, Wu Li, Wen-Xu Wang

**Author notes:** Corresponding authors (W.L.); (W.X.W.).

## Abstract

Neuroplasticity supports learning, development, and adaptive behavior, but also underlies maladaptive circuit remodeling in neuropsychiatric disorders. Although genetic disruptions of synaptic signaling are widely implicated in these conditions, the causal chain from molecular perturbation to systems-level dysfunction remains unclear. Here we introduce a multiscale computational framework—a parameter-free Boolean regulatory model of glutamatergic signaling—that captures the dynamic control of synaptic plasticity. The model generates emergent attractor states corresponding to long-term potentiation and depression and reveals how these states arise from underlying Hebbian principles. *In silico* gene-knockout simulations show that specific molecular perturbations destabilize these attractors, impairing synaptic plasticity and producing circuit-level alterations characteristic of diverse brain disorders. Remarkably, the model recapitulates the opposing developmental trajectories of autism spectrum disorder and schizophrenia, offering a mechanistic account of their divergent cortical phenotypes. This computationally tractable and generalizable framework quantitatively links genetic variation to synaptic instability and clinical severity, aligning with experimental and clinical observations. By moving beyond correlative associations, it establishes a causal scaffold for tracing how molecular disruptions propagate through synaptic regulatory networks to impair brain function, informing integrative diagnostics and mechanism-based therapeutic strategies.

## INTRODUCTION

Synaptic plasticity—the activity-dependent modification of synaptic strength—underlies learning, memory, and adaptive behavior (*Caroni et al., 2012; Citri and Malenka, 2008; Guskjolen and Cembrowski, 2023; Li, 2016; Martin et al., 2000; Neves et al., 2008*). Decades of work have characterized long-term potentiation (LTP) and long-term depression (LTD) at glutamatergic synapses (*Dan and Poo, 2006; Nicoll, 2017*), particularly in the hippocampus and cerebellum (*Bi and Poo, 1998; Bliss and Lømo, 1973; Dudek and Bear, 1992; Isaac et al., 1995*). These forms of plasticity depend on coordinated signaling through glutamatergic receptors, with AMPA receptors (AMPAR) trafficking at the postsynaptic membrane acting as a central effector mechanism (*Collingridge et al., 2004; Hammond, 2024; Hanley, 2018; Reiner and Levitz, 2018*).

Disruptions in these pathways are increasingly linked to a wide spectrum of neurological and psychiatric disorders (*Mattson et al., 2018; Price and Duman, 2020; Tartt et al., 2022*), including Alzheimer’s disease (AD), autism spectrum disorder (ASD), and schizophrenia (SCZ) (*Ebert and Greenberg, 2013; Ghosh and Giese, 2015; Kocahan and Dogan, 2017; Soler et al., 2018*). Genetic variations common in these conditions—such as single nucleotide polymorphisms and altered gene expression—alter synaptic signaling and plasticity, ultimately giving rise to circuit-level dysfunction and cognitive deficits. Despite abundant molecular and cellular evidence, it remains unclear how local perturbations propagate across biological scales to produce systems-level pathology. Computational models based on differential equations provide detailed descriptions of signaling components, but their reliance on numerous uncertain parameters limits interpretability and predictive power (*Barbano et al., 2007; Bell and Rangamani, 2023; Gallimore et al., 2016; Jȩdrzejewska-Szmek et al., 2017; Lee et al., 2024; Mäki-Marttunen et al., 2024; Mäki-Marttunen et al., 2020*). Given the nonlinear, combinatorial interactions among proteins, signaling cascades, and structural dynamics, a tractable systems-level approach is needed to understand how plasticity is regulated—and how its disruption contributes to disease.

Boolean network models, though operating on discrete states and time scales, provide an interpretable framework for examining complex biochemical regulatory networks. The robustness of biochemical reactions allows these discrete updates to approximate key regulatory functions. Although differential-equation models capture continuous dynamics, their parameter complexity can hinder interpretability and scalability (*Bell and Rangamani, 2023; Jȩdrzejewska-Szmek et al., 2017; Mäki-Marttunen et al., 2024; Mäki-Marttunen et al., 2020*), making Boolean modeling a practical alternative for dissecting regulatory logic. By emphasizing regulatory logic rather than kinetic detail, Boolean modeling isolates core control relationships in synaptic signaling and is well suited for assessing how molecular perturbations influence plasticity across molecular, synaptic, and circuit levels.

To address this need, we developed a multiscale framework that integrates molecular signaling with synaptic and circuit-level outcomes. At its core is a parameter-free Boolean network model of glutamatergic signaling, designed to capture the regulatory logic governing synaptic potentiation and depression. By exhaustively analyzing its dynamical landscape and systematically introducing *in silico* gene perturbations, this model reveals how molecular dysregulation destabilizes plasticity states and scales up to circuit abnormalities associated with neuropsychiatric disorders. This framework provides a mechanistic bridge from genetic variation to brain dysfunction and offers a computational foundation for understanding hierarchical disease progression and guiding therapeutics.

## RESULTS

### Boolean dynamics of synaptic plasticity

We simulated the dynamics of all 1,536 biologically plausible initial states to determine their trajectories toward stable attractors. Two representative examples are shown in Figure 3A, B, each starting with a distinct initial network state (T0). In both cases, the system consistently settled into a stable attractor state within a few time steps. Each attractor corresponds to a distinct biological outcome and falls into one of two categories: synaptic potentiation (P attractor; S_Endo_ = 0, S_SNAREs_ = 1; Figure 3A) or synaptic depression (D attractor; S_Endo_ = 1, S_SNAREs_ = 0; Figure 3B). A third attractor characterized by joint silence (S_Endo_ = 0, S_SNAREs_ = 0) also emerged, but we excluded it from further analysis due to its limited relevance to activity-dependent plasticity. Notably, no attractor displayed simultaneous activation of the two output nodes (Endo and SNARE), consistent with the mutual exclusivity of biological endocytosis and exocytosis pathways.

**Figure 1.**
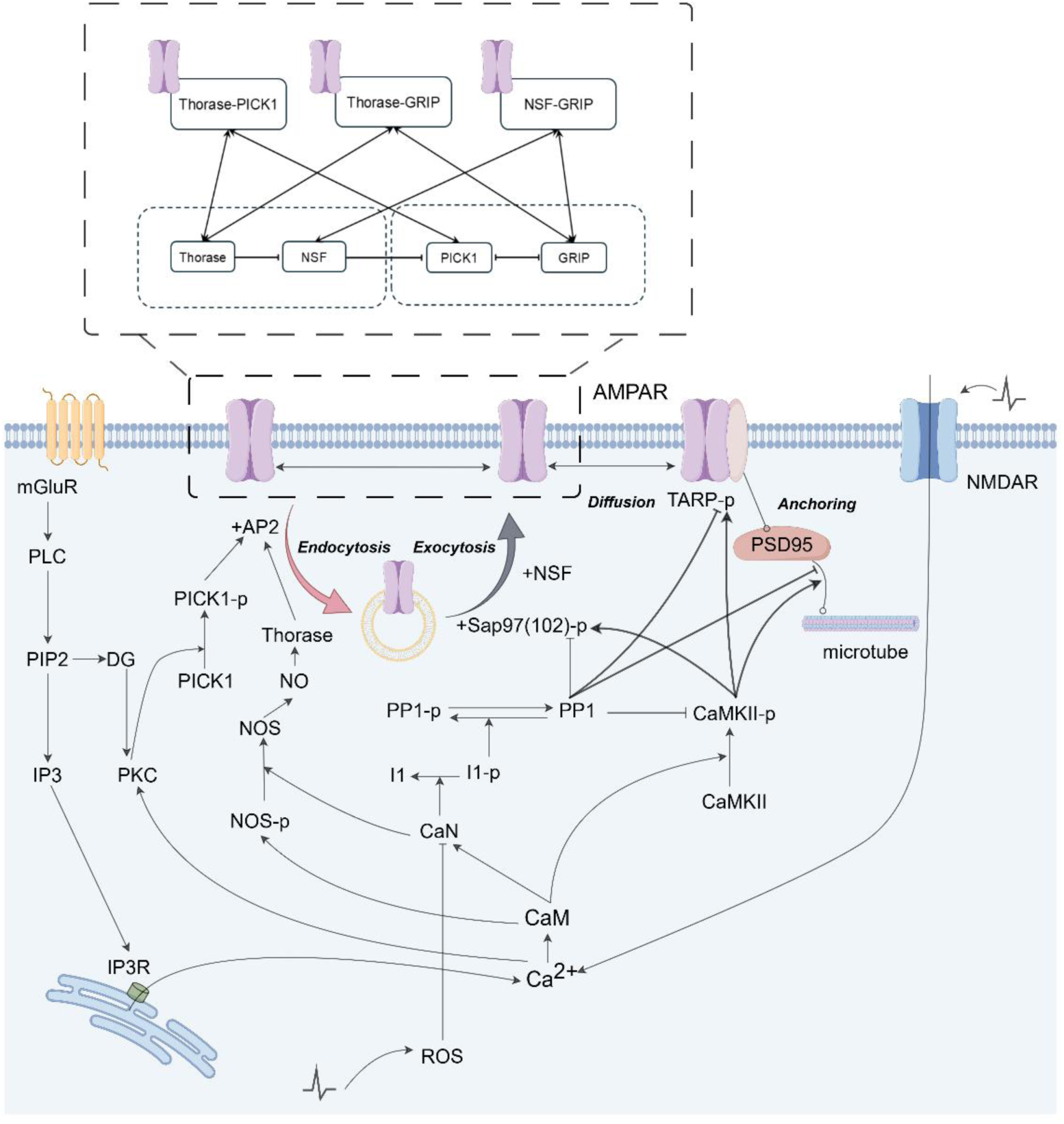
Postsynaptic signaling cascades and AMPAR trafficking. AMPAR trafficking is regulated by postsynaptic pathways triggered by mGluR and NMDAR activation, engaging both calmodulin- and PKC-mediated cascades. Calmodulin activates CaMKII and calcineurin (CaN), which play antagonistic roles: CaMKII promotes AMPAR stabilization at the synapse, whereas CaN facilitates dephosphorylation-dependent diffusion. Concurrently, PKC signaling modulates the phosphorylation of AMPAR subunits and their interacting proteins, critically influencing receptor diffusion and internalization. Within the postsynaptic density (PSD), AMPAR forms multiple binding states with auxiliary proteins that regulate receptor trafficking. Two pairs of proteins bind competitively to the GluR2 subunit: PICK1 and GRIP share one binding site, while Thorase and NSF occupy a separate site. NSF prevents PICK1 binding, whereas activated Thorase inhibits NSF interaction. The predominant PSD state is AMPAR–NSF–GRIP. Thorase displaces NSF, enabling a progressive transition into the AMPAR–Thorase–PICK1 complex, which is primed for endocytosis. Arrows indicate activation or catalysis; bar-headed lines indicate inhibition; circle-ended lines denote protein–protein interactions (e.g., between TARP and PSD95). Dashed boxes highlight proteins interacting with AMPAR that regulate receptor trafficking. See Supplemental table 1 for detailed biochemical interactions. Figure created using Figdraw.

**Figure 2.**
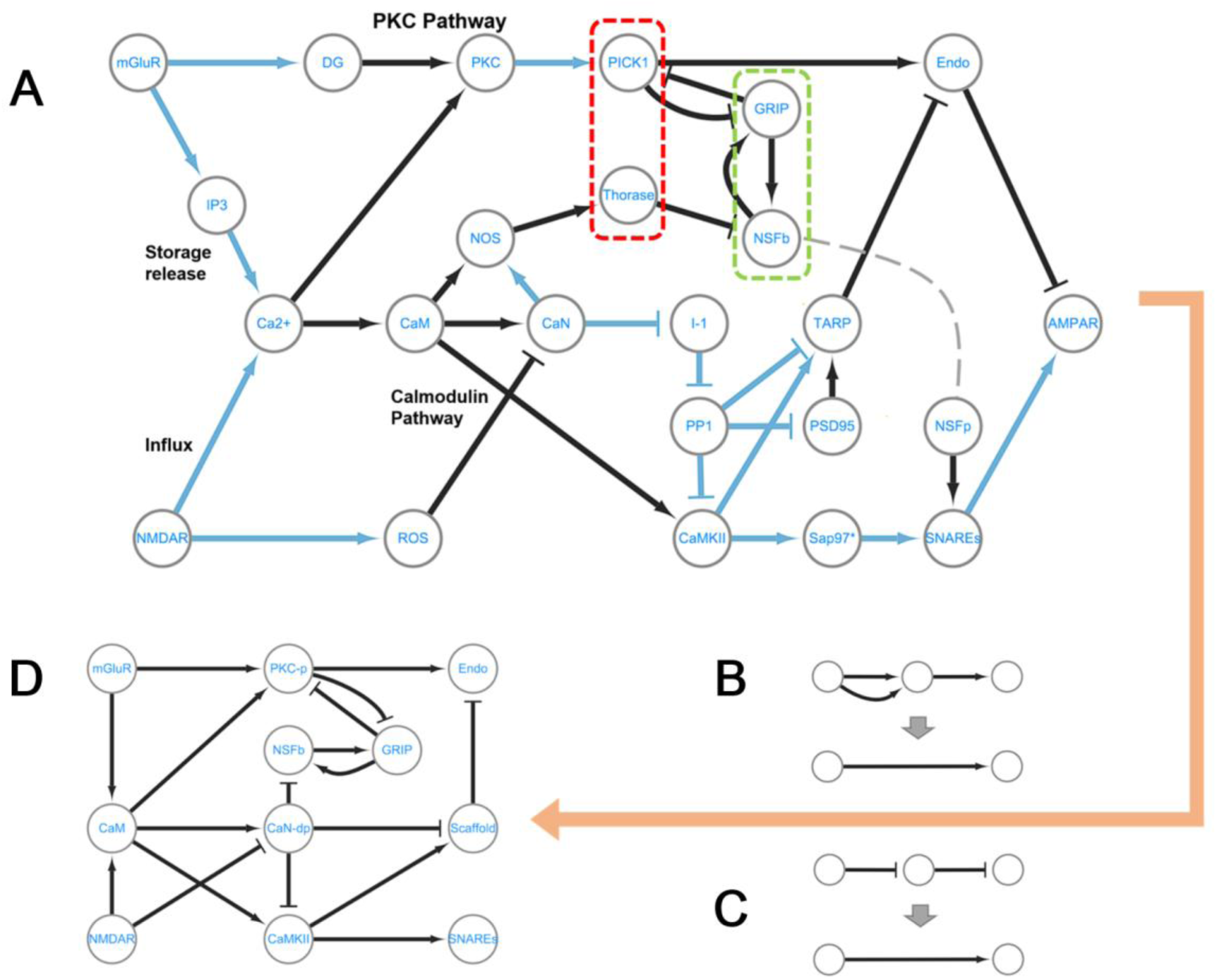
Synaptic plasticity regulatory network and its simplification. **(A)** The full Boolean network. It comprises 26 nodes and 39 edges, with edge color denoting two types of biochemical processes: black edges indicate binding reactions, while blue edges represent catalytic reactions. Each edge is further categorized by its regulatory effect: arrows denote activation, and T-shaped edges denote inhibition or blocking. The red and green dashed boxes highlight protein interactions with AMPAR, see also dashed box in Figure1. NSF is split into two functionally distinct nodes: NSFb represents AMPAR-bound NSF, while NSFp denotes the membrane-pool NSF, which assists in exocytosis (gray dashed curve). For better demonstration, Sap97/102 is omitted as Sap97*. For simplicity, proteins involved in exocytosis (e.g., Vamp, Syntaxin, and SNAP) are consolidated into a single node, SNAREs (lower right), while the protein complex responsible for endocytosis (including AP2, Dynamin, and Endophilin) is represented as the Endo node (upper right). Full node names and their Boolean functions are provided in Supplemental table 1. **(B**, **C)** Schematic of simplification rules. Multiple edges of the same type (activation or inhibition) between two nodes are merged into a single edge. If three nodes form a unidirectional activation chain (**B**), the middle node is merged with its downstream node. Similarly, for inhibition chains (**C**), the middle node is merged with the last node, and the entire inhibition chain is replaced by a single activation edge. **(D)** The simplified network. It consists of 11 nodes and 18 edges, with detailed edge and node merging listed in Supplemental table 2, and Boolean functions in Supplemental table 3. In particular, the major pathways from the full network are streamlined: CaN and PP1 are merged into CaN-dP; components of the PKC pathway are integrated into PKC-P; and TARP and PSD95 are combined into a single Scaffold node.

**Figure 3.**
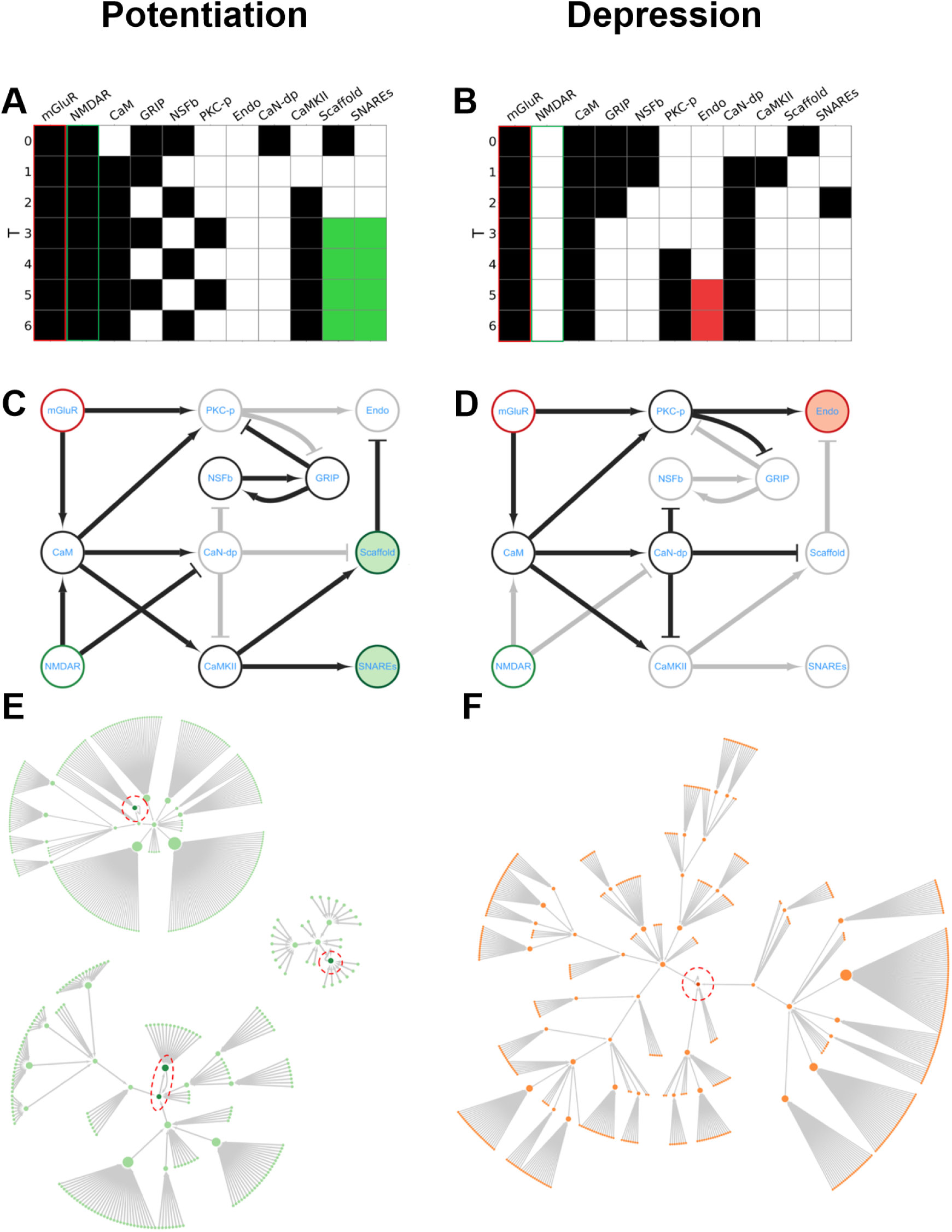
Evolution of the synaptic plasticity regulatory network in state space. **(A**, **B)** Examples of stepwise evolution of node states (T0 to T6), starting from distinct initial conditions (T0) and converging to the potentiation (P) and depression (D) attractors. Empty squares represent inactive state (0), and all filled squares indicate active state (1). The two input nodes, mGluR and NMDAR (left two columns), are outlined in red and green, respectively. The active states of the two output nodes, Endo and SNAREs, are highlighted with red and green filled squares, respectively. The P attractor corresponds to persistent activation of SNAREs (**A**), and the D attractor corresponds to persistent activation of Endo **(B)**. **(C, D)** Active (black) and inactive (gray) edges and nodes in the network when the system enters the attractor states shown in **(A)** and **(B)**. **(E, F)** Visualization of the P and D attractors and their respective attraction basins, shown in green (P) and orange (D). Attractors are enclosed in red dashed circles. The D attractor is a single fixed-point attractor (**F**), whereas the P attractor comprises three distinct configurations (**E**): two fixed-point attractors (top and right) and one period-two attractor (bottom), which oscillates between two states due to a self-sustaining loop involving NSFb and GRIP (see **C**). The size of each state node (representing a point in state space) is proportional to the number of neighboring states it connects to.

The active nodes and regulatory edges that define the P and D attractors in the Boolean network are highlighted in Figure 3C, D. A key feature of the P attractor is a self-sustaining loop involving NSFb and GRIP (Figure 3C), which maintains persistent activity even in the absence of inhibitory input to NSFb.

The attraction basins of the P and D attractors are visualized in Figure 3E, F. Each network state corresponds to a point in an 11-dimensional Boolean state space, encompassing 1,536 biologically plausible configurations. From any initial state, the system follows a deterministic trajectory—a sequence of discrete state transitions—toward one of the stable attractors. The set of all initial states that converge to a given attractor defines its attraction basin. The P and D attractors have distinct, non-overlapping attraction basins, meaning that each initial state leads exclusively to either synaptic potentiation or depression, but not both. This mutual exclusivity is biologically significant, as the simultaneous expression of these opposing mechanisms would disrupt proper synaptic modulation and, by extension, neuronal function.

Interestingly, one P attractor exhibited a period-two oscillation (Figure 3E), driven by the reciprocal interactions among NSFb, PKC-p, and GRIP in the simplified network (Figure 3C). To our knowledge, such an oscillatory P attractor has not been documented experimentally. This prediction offers a testable hypothesis regarding dynamic modes of molecular regulation during synaptic strengthening.

Overall, the Boolean dynamics recapitulate core principles of synaptic physiology. The emergence of the P attractor following NMDAR activation (Figure 3A, C, E) mirrors classical Hebbian learning (*Hebb, 2002; Lisman, 1989*). NMDAR activation requires coincident pre-synaptic glutamate release and post-synaptic depolarization—conditions met during synchronous neuronal firing. Through this coincidence-detection mechanism, the network reliably converges to the P attractor, thereby reinforcing synaptic strength in accordance with the classical rule: “fire together, wire together.”

### Plasticity impairment by node knockout

To investigate how molecular perturbations affect synaptic plasticity, we performed single-node knockout simulations on the full Boolean network, modeling the effects of individual gene mutations or protein dysfunction. For each perturbed network, we analyzed changes in the P and D attractors, including the number and type of stable states and the size of their attraction basins, compared with the intact network.

Note that knockouts were introduced prior to node and edge merging for network simplification. As a result, different knockouts can converge to the same simplified structure. From all possible single-node knockouts, we identified 11 distinct simplified residue networks. Figure 4 presents these networks alongside the intact network, showing their attractor states and quantifying the associated reduction in attraction basin size.

**Figure 4.**
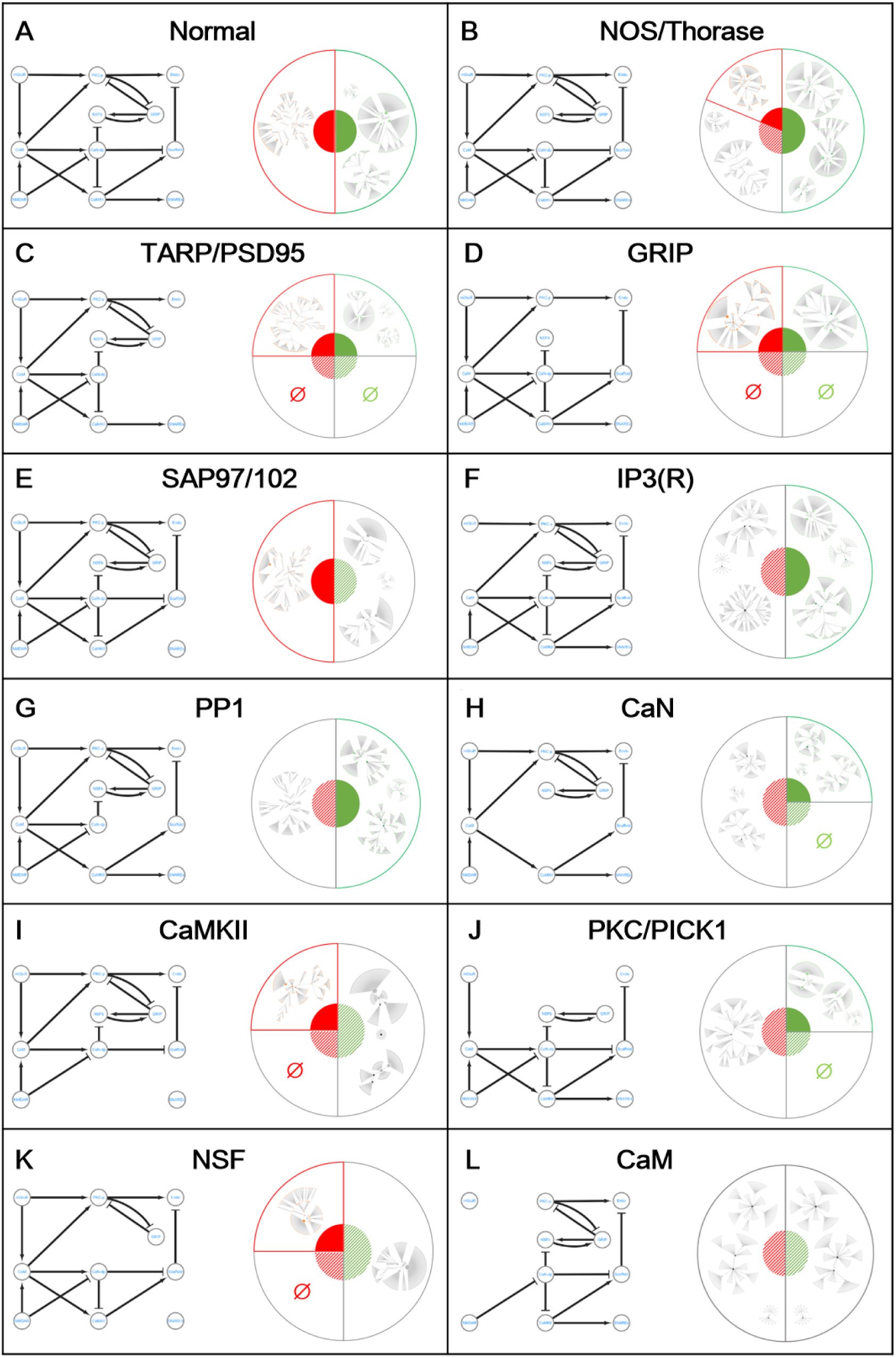
Residue networks, attractor states, and loss of attraction basins following single-node knockouts. **(A)** The intact network (left schematic) is shown together with the sizes of its potentiation (P) and depression (D) attraction basins, illustrated as half-pie charts filled in green and red, respectively (right). The corresponding attractor states and their full attraction basins—identical to those in Figure 3E, F except for a rearranged layout—are shown in the outer, unfilled pies. **(B**-**L)** Similar to (**A)**, these panels show the 11 residue networks resulting from single-node knockouts, ordered by increasing reductions in both P and D basin sizes relative to the intact network (combined shadowed green and red sectors), reflecting progressively greater impairment of synaptic plasticity. Two forms of basin disruption typically result from these knockouts: abnormal attractor substitution, where the native P and/or D attractor is replaced by a dysfunctional pattern (shown in the outer pies); and complete loss of the P and/or D basin, denoted by the null symbol (Ø).

The extent of attractor and attraction basin loss serves as a quantitative indicator of synaptic plasticity impairment. Larger loss of attraction basins reflects more severe plasticity damage. Node knockouts lead to varying degrees of impairment, measured as the fractional reduction in the attraction basin size (Equation 2). We classified the impairments into three categories: (1) mild damage (Figure 4B-D), where both potentiation and depression are preserved despite reduced basins; (2) severe damage (Figure 4E-K), where one function is entirely lost; and (3) lethal damage (Figure 4L), where both functions are lost, representing complete failure of synaptic plasticity. While direct experimental validation of these predictions remains challenging, our simulation results can be cross-referenced with empirical evidence from clinical research on various brain disorders associated with synaptic plasticity dysfunction.

### Plasticity impairment and brain disorder

Distinct brain disorders are characterized by specific structural and connectomic abnormalities, often stemming from opposing developmental processes. A prominent example is synaptic pruning: ASD has been linked to deficits in this process, potentially leading to over-connectivity, while SCZ is associated with excessive pruning—particularly in the frontal cortex—resulting in cortical thinning and under-connectivity. A multiscale causal framework is essential to explain how dysregulation of the same fundamental process at the synaptic level can propagate through neural circuits to produce such divergent clinical phenotypes.

During critical developmental periods, the precise timing of synaptic proliferation and pruning is fundamental for proper circuit formation and cognitive function. Disruptions in these plasticity mechanisms are implicated in the pathogenesis of neurodevelopmental disorders (*Forrest et al., 2018; Meredith, 2016*). We therefore hypothesize that the severity of brain dysfunction is positively correlated with and the degree of impairment in synaptic plasticity. To test this, we leveraged established genetic anomalies associated with brain diseases, as these provide a basis for aligning node knockout simulations in our Boolean model with specific neurological and psychiatric conditions.

Genetic studies have identified numerous risk loci with mutation variations across various brain disorders (Supplemental table 5). In our Boolean network model, node knockout operations simulate the functional impact of such mutations. To quantify disease severity, we adopted the WHO’s broad clinical classification, which categorizes disorders into two groups: diseases of the nervous system (DN) and mental, behavioral, or neurodevelopmental disorders (MBND), with DN generally representing more severe conditions (Supplemental table 4, 5). We simulated the knockout of nodes corresponding to these disease-associated loci and quantified the subsequent changes in the P and D attraction basins. Thus, within our framework, genetic anomalies serve as a bridge linking the model’s quantification of plasticity deficits to clinically defined disease severity, providing a causal test for our hypothesis.

Figure 5A illustrates the relationship between P- and D-basin loss and the severity of brain disorders. Each pie corresponds to one of the 11 networks from distinct single-node knockouts (see Figure 4). The pie coordinates indicate the degree of P- and D-basin loss, while the size and orange-to-blue ratio (DN vs. MBND cases) indicate relative prevalence. Notably, when P- and D-basin loss is smaller (closer to the lower left origin), DN cases are less frequent, as evidenced by smaller orange sectors. In contrast, severe P- and D-basin loss (close to the upper-right corner) correlates with a predominance of DN cases, where orange sectors exceed half of the pie. Figure 5B further demonstrates a significantly positive correlation between attraction-basin loss and disease severity (Pearson r = 0.7769, P = 0.0049, t-test), robustly supporting the concept of multiscale causality and validating our model’s predictive power.

**Figure 5.**
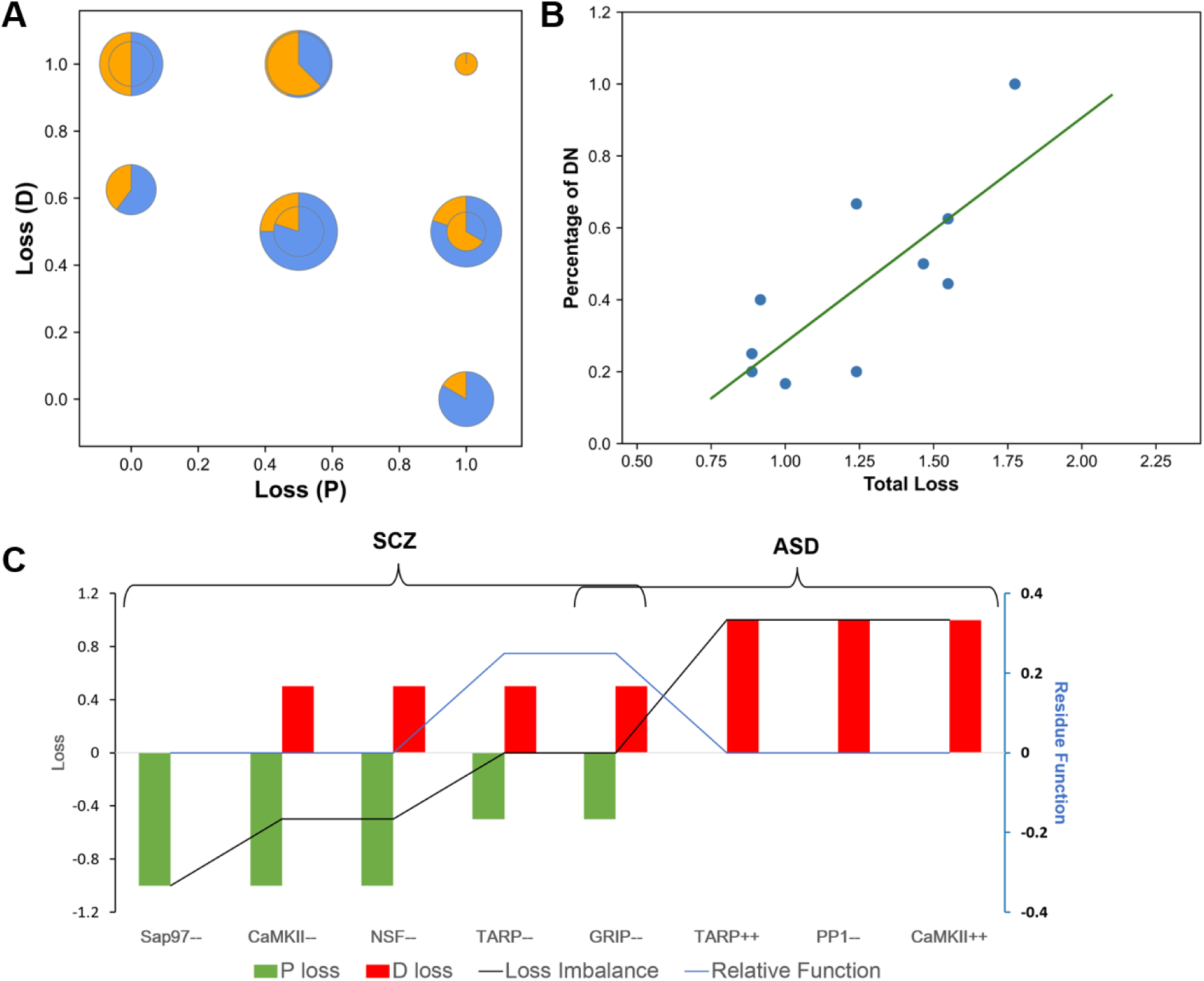
Relationship between attraction-basin loss and brain disease severity. **(A)** Loss of P- and D-attraction basins following single-node knockouts in the original network. Each pie chart corresponds to a residual network shown in Figure 4B-L. The pie center indicates the magnitude of attraction-basin loss (see Equation 2; overlapping centers reflect similar P- and D-basin losses in some residual networks). Pie size is proportional to the number of brain disease types associated with the corresponding gene deficit; orange and blue sectors denote DN and MBND cases, respectively. **(B)** Correlation between disease severity and attraction-basin loss (Pearson r = 0.7769, P = 0.0049, t-test). Each data point corresponds to one pie chart in **(A)**. Disease severity for each knockout is quantified as the percentage of the orange (DN) sector within the corresponding pie. For this specific analysis, total basin loss is defined as 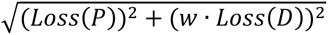, with weighting factor *w* = 1.465 (see S1 Fig for justification, which also shows that qualitatively similar results are obtained using the unweighted total loss). *Loss*(*P*) and *Loss*(*D*) denote the fractional losses of the P- and D-attractors, respectively (see Equation 2). Detailed disease information is provided in Supplemental table 5. **(C)** Modeling SCZ- and ASD-associated gene perturbations using Boolean network simulations. Clinical studies have identified 5 deficit-associated gene loci in SCZ and 4 in ASD, with GRIP implicated in both disorders. These loci are ordered by increasing difference between knockout-induced D-basin loss (red) and P-basin loss (green), as defined in Equation 2. The black curve shows the imbalance between depression and potentiation dynamics, calculated as *Loss*(*D*) − *Loss*(*P*). The blue curve shows a residue function capturing node-specific anomalies, computed as (*B*′_*D*_ · *B*′_*P*_)/(*B*_*D*_ · *B*_*P*_), where *B* and *B′* denote attraction-basin sizes in the intact and residue networks, respectively. Node annotations: (--) reduced expression or locational variation; (++) overexpression. Simulations of TARP and CaMKII overexpression are detailed in S2 Fig.

Neuroimaging and postmortem studies have demonstrated contrasting neuronal connectivity in ASD and SCZ. Adolescent ASD patients often exhibit a thicker cortex than healthy individuals, particularly in the frontal, temporal and parietal lobes (*Glantz and Lewis, 2001; Penzes et al., 2011*). In contrast, SCZ is characterizes by synaptic atrophy and reduced cortical thickness, especially in the frontal and temporal lobes, primary auditory cortex, and hippocampus (*Glantz and Lewis, 2001; Lewis and Levitt, 2002*). These morphological anomalies suggest that ASD and SCZ result from opposing synaptic plasticity dysfunctions. We investigated this by performing node knockout analysis on gene loci associated with each disorder.

Figure 5C quantifies the P- and D-basin loss (green and red bars, respectively) from knocking out genes linked to ASD and SCZ. The results reveal a differential impact: In ASD, synaptic depression impairment is more pronounced than potentiation impairment, whereas the opposite trend is observed in SCZ. This offers a molecular-level explanation for the distinct cortical abnormalities: impaired depression function in ASD leads to insufficient synaptic pruning and cortical thickening, while impaired potentiation function in SCZ promotes excessive pruning and cortical thinning.

Among the mutated genes, GRIP is notable for its implication in both ASD and SCZ. Its knockout leads to relatively mild, balanced impairments in both potentiation and depression (Figure 5C, GRIP--). We quantified this residual plasticity by assessing the network’s remaining functional capacity (Figure 5C, blue curve). This analysis suggests that GRIP mutations preserve more plasticity than other disease-associated genes. We speculate that as a shared comorbid factor in ASD and SCZ, GRIP dysfunction may establish a compensatory balance between potentiation and depression, resulting in comparatively milder symptoms.

## DISCUSSION

Our study provides a mechanistic framework for understanding how molecular-level disruptions in neuroplasticity can give rise to systems-level brain dysfunction. Although the importance of plasticity for learning, memory, and neurodevelopment is well established, the causal chains linking molecular perturbations to emergent circuit and behavioral abnormalities remain insufficiently defined. Here, we introduce a computational approach that explicitly traces how molecular regulation shapes synaptic dynamics and how these dynamics, when perturbed, contributes to pathological outcomes.

Focusing on glutamatergic synapses, we constructed a Boolean regulatory network that captures the core interactions governing AMPA receptor trafficking. Despite abstracting biochemical kinetics and simplifying regulatory details, the model retains essential causal logic and generates emergent attractor states corresponding to LTP and LTD. These attractors reflect stable receptor-trafficking motifs that enable synapses to transition between potentiated and depressed states in response to input structure.

The model primarily captures the induction phase of long-term potentiation and depression. In biological systems, sustained expression requires transcriptional regulation (e.g., CREB and NFAT activation downstream of CaMKII and CaN) and consolidation processes that depend on sleep and developmental stage. Because the transcriptional dynamics remain incompletely characterized and unfold on timescales beyond molecular signaling, our framework does not attempt to model long-term maintenance. Nonetheless, the stability of the P- and D-attractors is consistent with the anchoring effects of scaffolding proteins and activity-dependent receptor trafficking, suggesting that the regulatory logic captured here forms part of the mechanistic substrate upon which longer-term consolidation processes operate.

To examine how genetic dysregulation alters these synaptic states, we systematically knocked out nodes associated with genes implicated in neurological and psychiatric disorders. The model reveals that attractor stability is highly sensitive to specific molecular perturbations, and that the degree of attraction-basin reduction correlates positively with WHO disease severity scores. Notably, ASD- and SCZ-associated perturbations produce complementary impairments: ASD-linked dysregulations preferentially destabilize depression dynamics, consistent with reduced pruning and cortical thickening, whereas SCZ-linked dysregulations preferentially destabilize potentiation, consistent with excessive pruning and cortical thinning. These findings illustrate how distinct molecular disruptions can map onto divergent developmental trajectories and circuit phenotypes.

Differential-equation models provide valuable temporal precision, as exemplified by prior work modeling neuroplasticity mechanisms related to psychiatric disorders (*Bell and Rangamani, 2023; Gallimore et al., 2016; Jȩdrzejewska-Szmek et al., 2017; Mäki-Marttunen et al., 2024*). However, such models typically require extensive, and often uncertain, parameterization, limiting tractability and restricting analysis to biochemical reactions rather than enabling system-level metrics that link molecular perturbations to clinical outcomes. In contrast, Boolean models, constructed without free parameters, emphasize regulatory logic over kinetic details, allowing systematic interrogation of large networks while preserving key causal dependencies. This abstraction enables the identification of a generalizable system-level index—reduction of attraction-basin size—that predicts the impact of molecular dysregulation across disorders. This advantage reflects the robustness of biochemical signaling networks: the order and dependency of reactions often play a more decisive role than precise kinetic rates.

A central requirement for effective Boolean modeling is the accurate translation of molecular interactions into Boolean functions. If these functions fail to represent the underlying biology, system behavior will be mischaracterized. We attribute the limited success of previous Boolean approaches to incomplete reaction networks and oversimplified logic rules, rather than to intrinsic limitations of the Boolean formalism. Our construction, grounded in an extensive literature survey and the explicit incorporation of key signaling interactions, provides a more faithful representation that supports biologically interpretable inference.

This framework is extensible. Incorporating additional biological features—such as receptor subtypes (*Hell, 2016; Kennedy and Ehlers, 2006*), presynaptic signaling pathways, and glial contributions (*Eroglu and Barres, 2010; Sancho et al., 2021*)—would broaden its coverage of synaptic and circuit mechanisms. More generally, the value of such models lies in their ability to provide conceptual scaffolding for understanding how complex brain functions emerge from low-level molecular interactions. While data-driven approaches like deep learning have transformed tasks like protein structure prediction, modeling emergent regulatory logic remains an open challenge requiring both biologically grounded abstractions and experimental data.

In summary, by tracing the causal cascades from molecular dysregulation to systems-level pathology, this work introduces a generalizable platform for probing the hierarchical origins of brain disorders. The reduction of attraction basins provides a simple and quantitative marker for assessing how genetic or molecular perturbations influence disease risk—a model-based metric that, to our knowledge, has not been previously available. Future experimental and clinical work can further test and refine this metric, advancing the development of integrative diagnostics and mechanism-based therapeutic strategies.

## METHODS AND MATERIALS

### Molecular regulatory network of synaptic plasticity

At the molecular level, synaptic plasticity is implemented primarily through the regulation of AMPAR trafficking. We synthesized experimental findings (*Collingridge et al., 2004; Hanley, 2018; Huganir and Nicoll, 2013; Lisman et al., 2002; Sheng and Kim, 2002; Shepherd and Huganir, 2007; Song and Huganir, 2002*); see also Supplementary References 1-198) to construct a postsynaptic regulatory network that captures the core biochemical interactions controlling this process (Figure 1). The full network comprises 26 nodes and 39 directed edges representing key receptors, signaling proteins, and molecule complexes, along with their experimentally validated regulatory interactions (Figure 2A). Edges fall into two categories: binding interactions, which establish complexes that activate or inhibit downstream targets; and catalytic regulations, which reflect enzymatic effects on downstream activation states.

The network incorporates the major glutamate receptor classes: AMPAR, NMDAR, and mGluR. AMPAR functions as the principal output, while NMDAR and mGluR serve as input sources modulating intracellular signaling cascades (Figure 2A). Two canonical pathways (*Reiner and Levitz, 2018*) are embedded: the protein kinase C (PKC) cascade, initiated by mGluR activation (i.e., the mGluR–PKC pathway); and the Ca²⁺/calmodulin-CaMKII pathway, activated by calcium influx through NMDARs or release from endoplasmic reticulum (ER) stores. Additionally, the network includes more recently described mechanisms, such as Thorase-mediated AMPAR internalization via antagonism of N-ethylmaleimide-sensitive fusion protein (NSF), facilitated by nitric oxide. All interactions are experimentally substantiated (Supplemental table 1).

### Boolean formalization and network dynamics

We developed a Boolean network model to delineate the system’s core control principles without relying on fitted parameters. In this framework, each node adopts a binary state (active or inactive) and is updated by logical rules derived entirely from experimental evidence (Supplemental table 1). Consequently, the model’s dynamics are determined solely by prior biological knowledge, eliminating the need for parameter estimation and minimizing subjective assumptions.

We define the Boolean functions of each node based on biochemical reactions and molecular interactions identified in experimental studies. Since all biochemical processes in the regulatory network operate on comparable timescale, time is discretized and incorporated into the Boolean framework. The functional logic for nodes with different numbers of inputs is outlined below (a complete list of Boolean functions and their corresponding biochemical mechanisms are provided in Supplemental table 1):

1. Boolean function structure: Each node’s Boolean function specifies how its input signals jointly determine its state. These logical relationships reflect experimentally established biochemical interactions.
2. Single-input nodes: If the input activates the node, the output follows the input directly (e.g., *S*_*DG*_(*t* + 1) = *S*_*mGluR*_(*t*)). If the input inhibits the node, the output follows a NOT logic (e.g., *S*_*PP*1_(*t* + 1) = NOT *S*_*I*1_(*t*)).
3. Two-input nodes: If both excitatory inputs are required for activation, the relationship follows an AND logic (e.g., 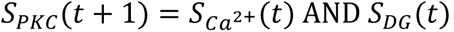). If either excitatory input alone is sufficient, the relationship follows an OR logic (e.g., 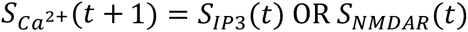). If one input is excitatory and the other inhibitory, and activation requires the presence of the excitatory input and the absence of the inhibitory input, the function follows an AND NOT logic (e.g., *S*_*Sap*97/102_(*t* + 1) = *S*_*CaMKII*_(*t*) AND NOT *S*_*PP*1_(*t*)).
4. Three-input nodes: Only the NSFb and TARP nodes have three inputs. Both receive two excitatory and one inhibitory input, but their Boolean functions differ. See Supplemental table 1 for specific logic definitions.

For any node *i*, its state *s*_*i*_(*t*) at time *t* is determined by its Boolean function and the states of its upstream inputs. The global state of the network at time *t*, denoted *S*(*t*), is defined as the collection of all node states:

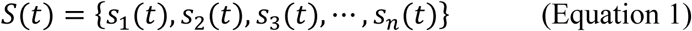

The set of all possible global states constitutes the network’s state space. In a convergent Boolean system, any initial state eventually evolves into a stable configuration known as an attractor. There are two types of attractors: fixed-point attractors, in which the network settles into a single, steady-state configuration; and limit-cycle attractors, in which the system enters a periodic loop, cycling through a finite sequence of states. Once an attractor is reached, the system’s global state *S*(*t*) repeatedly revisits one or more previously encountered configurations. By enumerating all possible initial states, we derive the full set of state-space trajectories and their corresponding attractors. The number of initial states that converge to a given attractor defines the size of its basin of attraction. In deterministic Boolean networks, each state belongs to one and only one attractor, ensuring that the state space is fully partitioned into non-overlapping attraction basins.

The model’s 26 nodes generate a state space of 2^26^ (∼67 million) configurations, making exhaustive analysis computationally intractable. To overcome this, we applied a biologically informed simplification that preserves essential network dynamics while enabling tractable analysis.

### Simplified Boolean network and functions

To reduce computational complexity while preserving essential regulatory logic, we systematically simplified the original molecular network under the constraint that its key dynamical features—specifically, the identity and stability of attractor states—remain intact. We applied two transformations (Figure 2B, C): First, node merging consolidated sequential activation chains into single composite units. Second, edge merging replaced multi-step or redundant interactions (e.g., double inhibition) with their net regulatory effect.

The full set of simplifications is detailed in Supplemental table 2. After simplification, the network is reduced to 12 nodes, each representing either a single molecular component or a composite functional unit. Critically, the network’s input structure is preserved: mGluR and NMDAR remain the only input nodes. AMPAR, the output node in the original network, is jointly regulated by two opposing intermediates: Endo (endocytosis machinery) and SNAREs (vesicular fusion proteins). To capture this antagonistic control more transparently, we omitted AMPAR in the simplified network and treated Endo and SNAREs as dual outputs, yielding a final network with 11 Boolean nodes and 18 directed edges (Figure 2D).

The reduced network has a theoretical state space of 2^11^ = 2,048 Boolean configurations. However, we restricted simulations to biologically feasible input states. For instance, NMDAR activation (*S*_NMDAR_ = 1) requires glutamate release and therefore co-occurring mGluRs activation (S_mGluR_ = 1); thus, the contradictory condition S_mGluR_ = 0 and *S*_NMDAR_ = 1 was excluded. Imposing all such constraints eliminated 2^9^ implausible configurations, yielding a final space of 1,536 biologically valid states.

This simplified Boolean network provides a computationally tractable platform for exhaustive simulations of synaptic plasticity dynamics. It retains the core regulatory logic of the full molecular network while allowing systematic investigation of how molecular perturbations propagate through the network to influence stable potentiation and depression states.

### Attractor damage and quantification of synaptic plasticity impairment

In the Boolean network’s state space, each attractor is associated with a basin of attraction comprising all initial states whose trajectories ultimately converge to that attractor. These basins can be visualized as tree-like structures, with multiple trajectories merging as they approach the same stable state. We quantify basin size as the total number of states that flow into a given attractor, and denote the basin sizes of the potentiation and depression attractors as *B*_*P*_ and *B*_*D*_, respectively.

To quantify the degree of synaptic plasticity impairment, we define the fractional loss of an attraction basin as

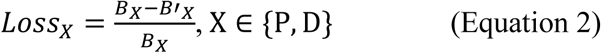

where *B*_*X*_ is the basin size of the potentiation (P) or depression (D) attractor in the intact network, and *B*′_*X*_ is the corresponding basin size in the perturbed (residual) network. Larger values of *Loss*_*X*_ indicate greater impairment of the corresponding form of synaptic plasticity.

### Psychiatric data and WHO classification

According to the World Health Organization (WHO), brain disorders are classified into two primary categories: (1) diseases of the nervous system (DN); (2) mental, behavioral, or neurodevelopmental disorders (MBND). In addition, certain genetic conditions such as Down syndrome and Fragile X syndrome, while not exclusively neurological, are associated with significant brain dysfunction. Due to the severity of their cognitive and behavioral manifestations, these disorders are treated as neurological diseases in this study (see Supplemental table 4 for details).

Based on a comprehensive literature review and accumulated data from four major databases (GWAS Catalog, ClinVar, GenCC and ClinGen), we compiled a list of genomic anomalies associated with brain diseases (summarized in Supplemental table 5). We integrated information from databases to summarize genes encoding nodal proteins and their associated, genetically validated brain diseases. Furthermore, we cross-referenced literature on brain disease risk loci with those focusing on genetically associated disorders, thereby summarizing the types of brain diseases associated with the plasticity regulation network. We categorized the genetic risk types (e.g., mutation, deletion, or SNPs). Given that disease-causing genetic variants are likely to impair the function of the original protein rather than enhance it, we pragmatically classified the identified genetic risks (from databases) as nodal impairments, considering those as enhancements only when there was clear evidence of overexpression. All protein subtypes in the brain with validated brain diseases are included in the analysis.

## ACKNOWLEDGMENTS AND DISCLOSURES

This study was supported by National Science and Technology Innovation 2030 Major Program 2022ZD0204600.

Z.Z.L., W.H.W., W.L. and W.X.W. conceptualized the study. Z.Z.L., W.X.W. and W.H.W. discussed the methodology. Z.Z.L. performed the investigation, data collection and simulation. W.H.W. helped visualization. Z.Z.L., W.X.W. and W.L. wrote the paper. W.H.W. conducted proof reading. The project was supervised by W.L. and W.X.W..

A previous version of this article was posted on bioRxiv as a preprint (https://www.biorxiv.org/content/10.1101/2025.04.07.647510v1).

The authors report no biomedical financial interests or potential conflicts of interest.

## REFERENCES

Barbano, P.E., Spivak, M., Flajolet, M., Nairn, A.C., Greengard, P., and Greengard, L. (2007) A mathematical tool for exploring the dynamics of biological networks Proceedings of the National academy of Sciences of the United States of America 104: 19169–19174.

Bell, M.K., and Rangamani, P. (2023) Crosstalk between biochemical signalling network architecture and trafficking governs AMPAR dynamics in synaptic plasticity Journal of Physiology 601: 3377–3402.

Bi, G.-q., and Poo, M.-m. (1998) Synaptic modifications in cultured hippocampal neurons: dependence on spike timing, synaptic strength, and postsynaptic cell type Journal of Neuroscience 18: 10464–10472.

Bliss, T.V.P., and Lømo, T. (1973) Long-lasting potentiation of synaptic transmission in the dentate area of the anaesthetized rabbit following stimulation of the perforant path Journal of Physiology 232: 331–356.

Caroni, P., Donato, F., and Muller, D. (2012) Structural plasticity upon learning: regulation and functions Nature Reviews Neuroscience 13: 478–490.

Citri, A., and Malenka, R.C. (2008) Synaptic plasticity: multiple forms, functions, and mechanisms Neuropsychopharmacology 33: 18–41.

Collingridge, G.L., Isaac, J.T., and Wang, Y.T. (2004) Receptor trafficking and synaptic plasticity Nature Reviews Neuroscience 5: 952–962.

Dan, Y., and Poo, M.-M. (2006) Spike timing-dependent plasticity: from synapse to perception Physiological Reviews 86: 1033–1048.

Dudek, S.M., and Bear, M.F. (1992) Homosynaptic long-term depression in area CA1 of hippocampus and effects of N-methyl-D-aspartate receptor blockade Proceedings of the National academy of Sciences of the United States of America 89: 4363–4367.

Ebert, D.H., and Greenberg, M.E. (2013) Activity-dependent neuronal signalling and autism spectrum disorder Nature 493: 327–337.

Eroglu, C., and Barres, B.A. (2010) Regulation of synaptic connectivity by glia Nature 468: 223–231.

Forrest, M.P., Parnell, E., and Penzes, P. (2018) Dendritic structural plasticity and neuropsychiatric disease Nature Reviews Neuroscience 19: 215–234.

Gallimore, A.R., Aricescu, A.R., Yuzaki, M., and Calinescu, R. (2016) A computational model for the AMPA receptor phosphorylation master switch regulating cerebellar long-term depression PLoS Computational Biology 12: e1004664.

Ghosh, A., and Giese, K.P. (2015) Calcium/calmodulin-dependent kinase II and Alzheimer’s disease Molecular Brain 8: 78.

Glantz, L.A., and Lewis, D.A. (2001) Dendritic spine density in schizophrenia and depression Archives of General Psychiatry 58: 203–203.

Guskjolen, A., and Cembrowski, M.S. (2023) Engram neurons: Encoding, consolidation, retrieval, and forgetting of memory Molecular Psychiatry 28: 3207–3219.

Hammond, C. (2024) Cellular and Molecular Neurophysiology, 5 edn (Academic Press).

Hanley, J.G. (2018) The regulation of AMPA receptor endocytosis by dynamic protein-protein interactions Frontiers in Cellular Neuroscience 12: 362.

Hebb, D.O. (2002) The Organization of Behavior, 1st edn (New York: Psychology Press).

Hell, J.W. (2016) How Ca2+-permeable AMPA receptors, the kinase PKA, and the phosphatase PP2B are intertwined in synaptic LTP and LTD Science Signaling 9: pe2–pe2.

Huganir, Richard L., and Nicoll, Roger A. (2013) AMPARs and synaptic plasticity: the last 25 years Neuron 80: 704–717.

Isaac, J.T.R., Nicoll, R.A., and Malenka, R.C. (1995) Evidence for silent synapses: implications for the expression of LTP Neuron 15: 427–434.

Jȩdrzejewska-Szmek, J., Luczak, V., Abel, T., and Blackwell, K.T. (2017) β-adrenergic signaling broadly contributes to LTP induction PLoS Computational Biology 13: e1005657.

Kennedy, M.J., and Ehlers, M.D. (2006) Organelles and trafficking machinery for postsynaptic plasticity Annual Review of Neuroscience 29: 325–362.

Kocahan, S., and Dogan, Z. (2017) Mechanisms of Alzheimer’s disease pathogenesis and prevention: the brain, neural pathology, N-methyl-D-aspartate receptors, tau protein and other risk factors Clinical Psychopharmacology and Neuroscience 15: 1–8.

Lee, C.T., Bell, M., Bonilla-Quintana, M., and Rangamani, P. (2024) Biophysical modeling of synaptic plasticity Annual Review of Biophysics 53: 397–426.

Lewis, D.A., and Levitt, P. (2002) Schizophrenia as a disorder of neurodevelopment Annual Review of Neuroscience 25: 409–432.

Li, W. (2016) Perceptual learning: use-dependent cortical plasticity Annual Review of Vision Science 2: 109–130.

Lisman, J. (1989) A mechanism for the Hebb and the anti-Hebb processes underlying learning and memory Proceedings of the National academy of Sciences of the United States of America 86: 9574–9578.

Lisman, J., Schulman, H., and Cline, H. (2002) The molecular basis of CaMKII function in synaptic and behavioural memory Nature Reviews Neuroscience 3: 175–190.

Mäki-Marttunen, T., Blackwell, K.T., Akkouh, I., Shadrin, A., Valstad, M., Elvsåshagen, T., Linne, M.-L., Djurovic, S., Einevoll, G.T., and Andreassen, O.A. (2024) Genetic mechanisms for impaired synaptic plasticity in schizophrenia revealed by computational modeling Proceedings of the National academy of Sciences of the United States of America 121: e2312511121.

Mäki-Marttunen, T., Iannella, N., Edwards, A.G., Einevoll, G.T., and Blackwell, K.T. (2020) A unified computational model for cortical post-synaptic plasticity eLife 9: e55714.

Martin, S.J., Grimwood, P.D., and Morris, R.G.M. (2000) Synaptic plasticity and memory: an evaluation of the hypothesis Annual Review of Neuroscience 23: 649–711.

Mattson, M.P., Moehl, K., Ghena, N., Schmaedick, M., and Cheng, A. (2018) Intermittent metabolic switching, neuroplasticity and brain health Nature Reviews Neuroscience 19: 81–94.

Meredith, R.M. (2016). Critical periods and neurodevelopmental brain disorders In Environmental Experience and Plasticity of the Developing Brain, A. Sale, ed. (John Wiley & Sons), pp. 73–98.

Neves, G., Cooke, S.F., and Bliss, T.V.P. (2008) Synaptic plasticity, memory and the hippocampus: a neural network approach to causality Nature Reviews Neuroscience 9: 65–75.

Nicoll, R.A. (2017) A brief history of long-term potentiation Neuron 93: 281–290.

Penzes, P., Cahill, M.E., Jones, K.A., VanLeeuwen, J.-E., and Woolfrey, K.M. (2011) Dendritic spine pathology in neuropsychiatric disorders Nature Neuroscience 14: 285–293.

Price, R.B., and Duman, R. (2020) Neuroplasticity in cognitive and psychological mechanisms of depression: an integrative model Molecular Psychiatry 25: 530–543.

Reiner, A., and Levitz, J. (2018) Glutamatergic signaling in the central nervous system: ionotropic and metabotropic receptors in concert Neuron 98: 1080–1098.

Sancho, L., Contreras, M., and Allen, N.J. (2021) Glia as sculptors of synaptic plasticity Neuroscience Research 167: 17–29.

Sheng, M., and Kim, M.J. (2002) Postsynaptic signaling and plasticity mechanisms Science 298: 776–780.

Shepherd, J.D., and Huganir, R.L. (2007) The cell biology of synaptic plasticity: AMPA receptor trafficking Annual Review of Cell and Developmental Biology 23: 613–643.

Soler, J., Fananas, L., Parellada, M., Krebs, M.-O., Rouleau, G.A., and Fatjo-Vilas, M. (2018) Genetic variability in scaffolding proteins and risk for schizophrenia and autism-spectrum disorders: a systematic review Journal of Psychiatry & Neuroscience 43: 223–244.

Song, I., and Huganir, R.L. (2002) Regulation of AMPA receptors during synaptic plasticity Trends in Neurosciences 25: 578–588.

Tartt, A.N., Mariani, M.B., Hen, R., Mann, J.J., and Boldrini, M. (2022) Dysregulation of adult hippocampal neuroplasticity in major depression: pathogenesis and therapeutic implications Molecular Psychiatry 27: 2689–2699.

