## Supplementary Materials for "A computational framework linking molecular regulation, synaptic plasticity, and brain disorders"

**This file includes:**

Supplementary Figure 1 to 2
Supplementary Tables 1 to 5
Supplementary References

### Supplementary figures

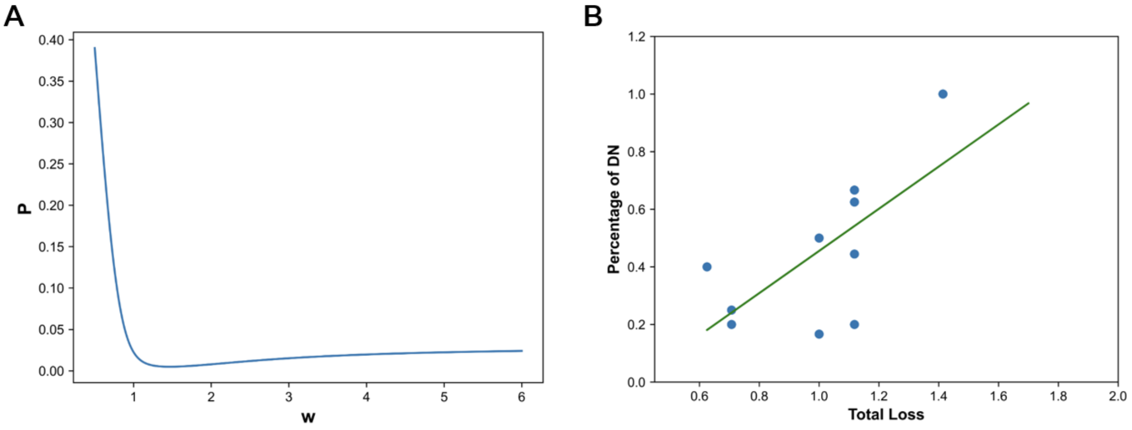

**S1 Fig. Optimization of the weighted total loss and comparison with the unweighted metric for computing attraction-basin loss.**

The weighting factor used in Figure 5B accounts for the fact that damage to the potentiation (P) and depression (D) attractors may contribute unequally to the prevalence of “diseases of the nervous system” (DN; see Table S4). This asymmetry likely arises because synaptic potentiation is supported by auxiliary pathways—such as PKA and Akt—in addition to CaMKII, whereas no alternative pathways are known for CaN- and PKC-mediated regulation of AMPARs. To address this imbalance, a weighting factor *w* was introduced in the total basin loss calculation used specifically in Figure 5B, where $Total\ Loss = \sqrt{(Loss(P))^2 + (w \cdot Loss(D))^2}$, with $Loss(P)$ and $Loss(D)$ denoting the fractional losses of the P- and D-attractors, respectively (see Equation 2).

**(A)** Optimization of the weighting factor. Linear regression was performed between the weighted total basin loss and DN prevalence for varying *w*, and the resulting P-values are plotted as a function of *w*. The minimum P value (P = 0.0049) occurs at *w* = 1.465. For clarity, only *w* = 0.5 to 6 is shown.

**(B)** Correlation between brain disease severity and the unweighted total basin loss ($\sqrt{(Loss_P)^2 + (Loss_D)^2}$; Pearson r = 0.675, P = 0.0226, t-test), analogous to Figure 5B but computed without the weighting factor (i.e., *w* = 1).

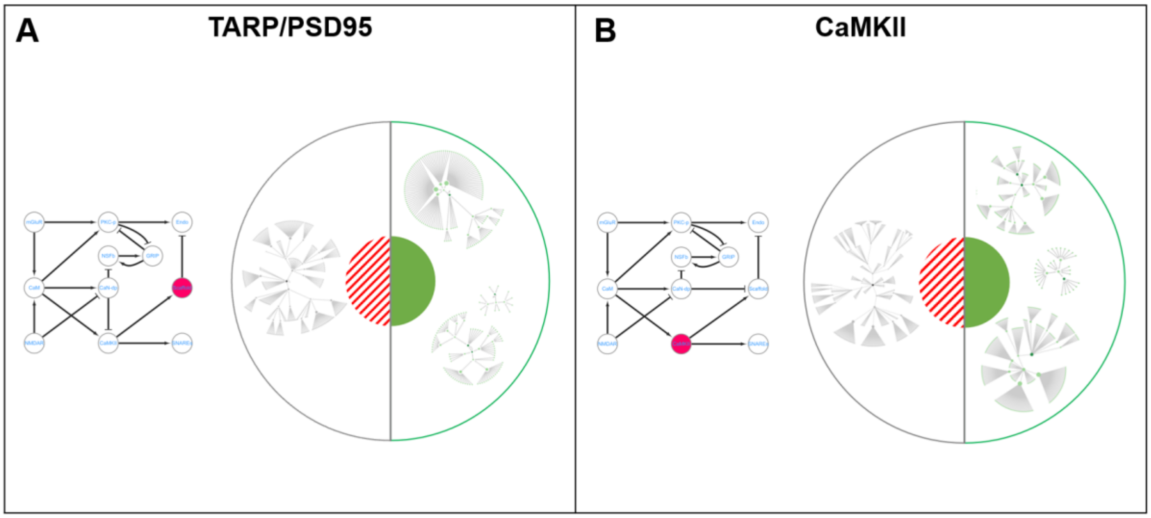

**S2 Fig. Attractors and attraction basins under node overexpression.**
**(A)** Left: Simplified network with overexpression of the Scaffold node (representing TARP/PSD95), which is highlighted in red. The resulting attractor states and their attraction basins are visualized using the same scheme as in Figure 4.
**(B)** Same as in **(A)**, but for overexpression of CaMKII.

Note:
In our comparative analysis of ASD and SCZ (Figure 5C), we incorporated the overexpression of PSD95, TARP, and CaMKII based on empirical findings. In the simplified network, TARP and PSD95 are merged into a single Scaffold node, resulting in two overexpression targets: Scaffold and CaMKII. The original Boolean function for Scaffold, $S_{Scaffold}(t+1) = S_{CaMKII}(t)\ \mathrm{OR\ NOT}\ S_{PP1}(t)$ is replaced with $S_{Scaffold}(t+1) = 1$. This simulates constitutive overexpression due to the absence of inhibitory regulation. As for CaMKII, $S_{CaMKII}(t+1) = S_{CaM}(t)\ \mathrm{AND\ NOT}\ S_{PP1}(t)$ is modified to $S_{CaMKII}(t+1) = S_{CaM}(t)$. This reflects a loss of PP1-mediated inhibition, resulting in persistent CaMKII activation. These modifications preserve the structure of the network while adjusting the logical dynamics to reflect gene overexpression. The simulation procedure remains identical to that of the intact simplified network.

**Supplementary Table 1. Boolean functions of nodes and their biochemical relevance.**

| Node centered at biochemical reactions | Biochemical reactions | | Boolean functions |
|---|---|---|---|
| Diacylglycerol/ Inositol triphosphate (DG/IP3) | mGluR (subtypes 1/5) binds glutamate and activates G-protein(1);<br>G-protein then activates downstream PLC and decomposes PIP2(1, 2) | $mGluR + Glu \rightarrow PLC*$<br>$PIP2 \xrightarrow{PLC*} DG + IP3$ | $S_{DG}(t+1) = S_{mGluR}(t)$,<br><br>$S_{IP3}(t+1) = S_{mGluR}(t)$ |
| Calcium (Ca2+) | IP3 binds IP3 receptors (IP3R) on Endoplasmic Reticulum(ER) (2);<br>Calcium stored in ER released through opened IP3R (3);<br>Calcium influx through opened NMDAR (4). | $IP3R \xrightarrow{IP3} IP3R*$<br>$Ca^{2+}(store) \xrightarrow{IP3R*} Ca^{2+}$<br>$Ca^{2+}(Out) \xrightarrow{NMDAR*} Ca^{2+}$ | $S_{Ca^{2+}}(t+1) = S_{IP3}(t)\text{OR}\ S_{NMDAR}(t)$ |
| Reactive oxygen species (ROS) | ROS is generated when post-synaptic neuron fires (5) ;<br>Here post-synaptic neuron firing is necessary for NMDAR activation (4);<br>So, ROS is of the same state as NMDAR. | Logically，during neuron firing (Depolarization):<br>$Mitochondira \rightarrow ROS$<br>$NMDAR \xrightarrow{Glu+\text{Depolarization}} NMDAR*$<br><br>Thus, logically:<br>$S_{ROS} = 1$ when $S_{NMDAR} = 1$ | $S_{ROS}(t+1) = S_{NMDAR}(t)$ |
| Calmodulin (CaM) | CaM changes to an active form through binding $Ca^{2+}$ (4, 6). | $CaM + Ca^{2+} \rightarrow CaM*$ | $S_{CaM}(t+1) = S_{Ca^{2+}}(t)$ |
| Calcineurin (CaN) | The $Ca^{2+}$ combined CaM activates CaN's catalytic activity (7, 8);<br>ROS inhibit CaN's catalytic activity (9-11). | $CaN + CaM* \rightarrow CaN*$<br><br>$CaN* \xrightarrow{ROS} CaN(inhibited)$ | $S_{CaN}(t+1) = S_{CaM}(t)$ AND NOT$S_{ROS}(t)$ |
| Protein Kinase C (PKC) | PKC has two binding domains C1 and C2, where DAG binds at C1 and $Ca^{2+}$ binds at C2.<br>PKC requires both for activation (12, 13). | $PKC + DG + Ca^{2+} \rightarrow PKC*$ | $S_{PKC}(t+1) = S_{Ca^{2+}}(t)$ AND $S_{DG}(t)$ |
| Protein Interacting with C Kinase 1 (PICK1) | PICK1 is phosphorylated by PKC to be active (14, 15);<br>Activated PICK1 binds AMPAR subunits GluA2/3 to trigger their phosphorylation and downstream reactions (15);<br>GRIP competes with PICK1 for the same AMPAR binding site (16). | $PICK1 \xrightarrow{PKC*} PICK1_p$<br><br>$AMPAR + PICK1_p \rightarrow AMPAR_{-PICK1_p}$<br><br>$AMPAR_{-PICK1_p} + GRIP \leftrightarrow AMPAR_{-GRIP} + PICK1_p$<br><br>$S_{PICK1} = 1$ represents generation of $AMPAR_{-PICK1_p}$ | $S_{PICK1}(t+1) = S_{PKC}(t)$ AND NOT$S_{GRIP}(t)$ |
| Glutamate Receptor- | PICK1 competes with GRIP for the same AMPAR binding site (16, 17). | $AMPAR + GRIP \rightarrow AMPAR_{-GRIP}$ | $S_{GRIP}(t+1) =$ NOT $S_{PICK1}(t)$ AND $S_{NSFb}(t)$ |

| | | | |
|---|---|---|---|
| Interacting Protein (GRIP) | NSF binds AMPAR to block PICK1's binding with AMPAR, so that facilitates GRIP's binding with AMPAR and its stabilization (18, 19). | $AMPAR_{-PICK1_p} + GRIP \leftrightarrow AMPAR_{-GRIP} + PICK1_p$<br><br>$AMPAR_{-PICK1_p} + GRIP + NSF \rightarrow AMPAR_{-NSF-GRIP} + PICK1_p$<br><br>$S_{GRIP} = 1$ represents generation of $AMPAR_{-GRIP}$ | |
| N-ethylmaleimide-Sensitive Fusion protein (NSF) | NSF is sufficient on the membrane, and it automatically binds AMPAR if it's not inhibited (20, 21). | See details in the left column | $S_{NSF}(t) = 1$ |
| NSF-bound (NSFb) | NSFb refers to AMPAR-bound NSF (20, 22).<br>Thorase competes with NSFb for the same AMPAR binding domain(23).<br>NSF is more likely to bind AMPAR with GRIP. | $AMPAR + NSF \rightarrow AMPAR_{-NSF}$<br><br>$AMPAR_{-GRIP} + NSF \rightarrow AMPAR_{-NSF-GRIP}$<br><br>$AMPAR_{-NSF(-GRIP)} + Thorase* \rightarrow AMPAR_{-Thorase*(-GRIP)}$ | $S_{NSFb}(t+1) = \text{NOT} S_{Thorase}(t)$ AND $S_{GRIP}(t)$ AND $S_{NSF}(t)$ |
| Thorase | Thorase is activated by Nitrogen Monoxide (NO), where NO is produced by NOS (24). | $Thorase \xrightarrow{NO} Thorase*$<br><br>$L - Arg \xrightarrow{NOS*_p} NO$ | $S_{Thorase}(t+1) = S_{NOS}(t)$ |
| Nitric Oxide Synthase (NOS) | NOS need to bind CaM and be dephosphorylated by CaN in order to produce NO (25). | $NOS + CaM* \rightarrow NOS*$<br><br>$NOS* \xrightarrow{CaN*} NOS*_p$<br><br>$S_{NOS} = 1$ for state $NOS*_p$ | $S_{NOS}(t+1) = S_{CaM}(t)$AND $S_{CaN}(t)$ |
| Inhibitor-1 (I-1) | The active I-1 or DARPP-32 for PP1 inhibition is dephosphorylated by CaN (26). | $I1_p \xrightarrow{CaN*} I1$<br><br>$S_{I1} = 1$ for state $I1_p$ | $S_{I1}(t+1) = \text{NOT } S_{CaN}(t)$ |
| Protein Phosphatase1 (PP1) | I-1 inhibits PP1's activity(27, 28). | $PP1 + I1_p \rightarrow I1 + PP1_p$<br><br>$S_{PP1} = 1$ for state $PP1$ | $S_{PP1}(t+1) = \text{NOT } S_{I1}(t)$ |
| Calcium-calmodulin (CaM)-dependent protein Kinase II (CaMKII) | CaMKII binds CaM to gain ability to phosphorylate substrates (29);<br>PP1 inhibit CaMKII's phosphorylation ability (30). | $CaMKII + CaM* \rightarrow CaMKII_p$<br><br>$CaMKII_p \xrightarrow{PP1} CaMKII$<br><br>$S_{CaMKII} = 1$ for state $CaMKII_p$ | $S_{CaMKII}(t+1) = S_{CaM}(t)$AND NOT $S_{PP1}(t)$ |
| Synapse-associated protein 97/102 (Sap97/102) | Sap97/102 are the main scaffolds of GluA1/A2's translocation, respectively (31-33).<br>Sap97/102 is activated by CaMKII's phosphorylation and deactivated by PP1's dephosphorylation (34, 35). | $Sap97 \xrightarrow{CaMKII_p} Sap97_p$<br><br>$Sap102 \xrightarrow{CaMKII_p} Sap102_p$<br><br>$Sap97_p \xrightarrow{PP1} Sap97$<br><br>$Sap102_p \xrightarrow{PP1} Sap102$ | $S_{Sap97/102}(t+1) = S_{CaMKII}(t)$ AND NOT $S_{PP1}(t)$ |

| | | | |
|---|---|---|---|
| Transmembrane AMPAR Regulatory Proteins (TARP) | Phosphorylated TARP implements its function by binding PSD95 (36, 37). TARP's phosphorylation is jointly determined by CaMKII and PP1. In the absence of both CaMKII and PP1, TARP's normal phosphorylation states are not affected. The presence of CaMKII ensures phosphorylation, regardless of PP1. Without CaMKII, the existence of PP1 dephosphorylate TARP, so that the necessary condition is not satisfied (29, 38, 39). | $AMPAR + TARP \rightarrow AMPAR_{-TARP}$<br><br>$AMPAR_{-TARP} \xrightarrow{CaMKII_p} AMPAR_{-TARP_p}$<br><br>$AMPAR_{-TARP_p} \xrightarrow{PP1} AMPAR_{-TARP}$<br><br>$AMPAR_{-TARP_p} + PSD95 * \rightarrow AMPAR_{-TARP_p}\ (Anchored)$<br><br>$S_{TARP}$=1 for state $AMPAR_{-TARP_p}\ (Anchored)$. | $S_{TARP}(t+1) = S_{CaMKII}(t)$ OR NOT $S_{PP1}(t)$ AND $S_{PSD95}(t)$ |
| Postsynaptic Density protein-95 (PSD95) | PSD95 is the main post-synaptic scaffold protein (37, 40).<br>In the absence of dephosphorylation from PP1, PSD95 always exists with $S_{PSD95} = 1$ (30, 41). | $PSD95 * \xrightarrow{PP1} PSD95$ | $S_{PSD95}(t+1) = 1\ AND\ NOT\ S_{PP1}(t)$ |
| Endo*(refers to Endocytosis protein complex) | Endo is a complex of necessary proteins for endocytosis (internalization), such as AP2, Dynamin, Endophilin, etc (42).<br>PICK1 binds both AP2 and AMPAR to form an endocytosis complex (43).<br>When binding to PSD95, TARPs inhibit AMPARs from lateral diffusion, hence inhibit endocytosis (38, 44). | $AMPAR + PICK1_p \xrightarrow{AP2} AMPAR_{-PICK1_p}\ (endocytosis)$<br><br>$AMPAR_{-TARP} + PICK1_p \xrightarrow{AP2} AMPAR_{-TARP-PICK1_p}\ (endocytosis)$<br><br>$AMPAR_{-TARP_p}\ (anchored)$ is not to enter endocytosis throw AP2 interaction.<br><br>$S_{Endo} = 1$ for all $AMPAR_{-X\ldots}\ (endocytosis)$ | $S_{Endo}(t+1) = S_{PICK1}(t)$AND NOT $S_{TARP}(t)$ |
| Soluble *N*-ethylmaleimide-sensitive factor attachment protein receptors (SNAREs) | AMPAR needs to bind phosphorylated Sap97/102 to be translocated and bind SNAREs (33, 35, 45, 46)； SNAREs need NSF for exocytosis (47). | $AMPAR(endosome) + Sap97_p \rightarrow AMPAR_{-Sap97_p}\ (endosome)$<br><br>$AMPAR_{-Sap97_p}\ (endosome) + SNAREs \xrightarrow{NSF} AMPAR(exocytosis)$<br><br>$S_{SNAREs} = 1$ for $AMPAR(exocytosis)$ | $S_{SNAREs}(t+1) = S_{\frac{Sap97}{102}}(t)$AND $S_{NSF}(t)$ |
| α-amino-3-hydroxy-5-methyl-4-isoxazoliopropionate receptor (AMPAR) | AMPARs regulation relies on insertion and internalization through exocytosis and endocytosis, respectively (48).<br>SNAREs is a necessary protein complex for AMPAR exocytosis (49) (insertion, $S_{AMPAR} = 1$). | $AMPAR(increment) = AMPAR(exocytosis) - AMPAR(endocytosis)$ | $S_{AMPAR}(t+1) = S_{SNAREs}(t)$ AND NOT $S_{Endo}(t$ |

| | |
|---|---|
| | Endo is a complex of necessary proteins for AMPAR endocytosis (49) (internalization, $S_{AMPAR} = 0$ ). $S_{SNAREs} = 0$ $and$ $S_{Endo} = 0$ are irrelevant to plasticity, so have no Boolean output. $S_{SNAREs} = 1$ $and$ $S_{Endo} = 1$ are biologically abnormal, so also have no Boolean output. |

Note:

1. References to biochemical reactions/interactions (middle column) are indicated by parenthesized numbers (see Supplementary References).
2. '*' refers to activation states for enzymes and proteins.
3. Subscript *p* refers to phosphorylation of proteins.
4. $AMPAR_{-X}$ refers to X protein binding to AMPAR.

**Supplementary Table 2. Pathway shortening in the original Boolean network.**

| Original nodes and edges | Nodes and edges after simplification |
|---|---|
| mGluR → DG → PKC → PICK1 | mGluR → PKC-p* |
| mGluR → IP3R → $Ca^{2+}$ → CaM | mGluR → CaM |
| → Ca2+ → CaM → | → CaM → |
| CaN ¬ I-1 ¬ PP1 ¬ CaMKII | CaN-dp* ¬ CaMKII |
| CaN-dp* → NOS→ Thorase ¬ NSFb | CaN-dp* ¬NSFb |
| NMDAR → ROS ¬ CaN | NMDAR ¬ CaN-dp* |
| CaMKII → Sap97 → SNAREs | CaMKII → SNAREs |
| PSD95 → TARP ¬ Endo | Scaffold* ¬ Endo |

Note:

1. Nodes marked with '*' derive from node merging.
2. '→'denotes activation
3. '¬' denotes inhibition.

**Supplementary Table 3. Boolean functions of nodes in the simplified Boolean network.**

<table>
<tr><th>Nodes name</th><th>Boolean function</th><th colspan="4">States detail</th></tr>
<tr><td rowspan="9">PKC-p</td><td rowspan="9">$S_{PKC_p}(t+1) =$<br>$S_{mGluR}(t)$ AND $S_{CaM}(t)$ AND NOT $S_{GRIP}(t)$</td><td>$S_{mGluR}(t)$</td><td>$S_{CaM}(t)$</td><td>$S_{GRIP}(t)$</td><td>$S_{PKC_p}(t+1)$</td></tr>
<tr><td>0</td><td>0</td><td>0</td><td>0</td></tr>
<tr><td>0</td><td>1</td><td>0</td><td>0</td></tr>
<tr><td>1</td><td>0</td><td>0</td><td>0</td></tr>
<tr><td>1</td><td>1</td><td>0</td><td>1</td></tr>
<tr><td>0</td><td>0</td><td>1</td><td>0</td></tr>
<tr><td>0</td><td>1</td><td>1</td><td>0</td></tr>
<tr><td>1</td><td>0</td><td>1</td><td>0</td></tr>
<tr><td>1</td><td>1</td><td>1</td><td>0</td></tr>
<tr><td rowspan="5">CaM</td><td rowspan="5">$S_{CaM}(t+1) =$<br>$S_{mGluR}(t)$ OR $S_{NMDAR}(t)$</td><td>$S_{mGluR}(t)$</td><td>$S_{NMDAR}(t)$</td><td colspan="2">$S_{CaM}(t+1)$</td></tr>
<tr><td>0</td><td>0</td><td colspan="2">0</td></tr>
<tr><td>0</td><td>1</td><td colspan="2">1</td></tr>
<tr><td>1</td><td>0</td><td colspan="2">1</td></tr>
<tr><td>1</td><td>1</td><td colspan="2">1</td></tr>
<tr><td rowspan="5">CaN-dP</td><td rowspan="5">$S_{CaN-dp}(t+1) =$<br>$S_{CaM}(t)$ AND NOT $S_{NMDAR}(t)$</td><td>$S_{CaM}(t)$</td><td>$S_{NMDAR}(t)$</td><td colspan="2">$S_{CaN-dp}(t+1)$</td></tr>
<tr><td>0</td><td>0</td><td colspan="2">0</td></tr>
<tr><td>0</td><td>1</td><td colspan="2">0</td></tr>
<tr><td>1</td><td>0</td><td colspan="2">1</td></tr>
<tr><td>1</td><td>1</td><td colspan="2">0</td></tr>
<tr><td rowspan="5">NSFb</td><td rowspan="5">$S_{NSFb}(t+1) =$<br>$S_{GRIP}(t)$ AND NOT $S_{CaN-dp}(t)$</td><td>$S_{GRIP}(t)$</td><td>$S_{CaN-dp}(t)$</td><td colspan="2">$S_{NSFb}(t+1)$</td></tr>
<tr><td>0</td><td>0</td><td colspan="2">0</td></tr>
<tr><td>0</td><td>1</td><td colspan="2">0</td></tr>
<tr><td>1</td><td>0</td><td colspan="2">1</td></tr>
<tr><td>1</td><td>1</td><td colspan="2">0</td></tr>
<tr><td rowspan="5">GRIP</td><td rowspan="5">$S_{GRIP}(t+1) =$<br>$S_{NSFb}(t)$ AND NOT $S_{PKC-p}(t)$</td><td>$S_{NSFb}(t)$</td><td>$S_{PKC-p}(t)$</td><td colspan="2">$S_{GRIP}(t+1)$</td></tr>
<tr><td>0</td><td>0</td><td colspan="2">0</td></tr>
<tr><td>0</td><td>1</td><td colspan="2">0</td></tr>
<tr><td>1</td><td>0</td><td colspan="2">1</td></tr>
<tr><td>1</td><td>1</td><td colspan="2">0</td></tr>
<tr><td rowspan="5">CaMKII</td><td rowspan="5">$S_{CaMKII}(t+1) =$<br>NOT $S_{CaN-dp}(t)$ AND $S_{CaM}(t)$</td><td>$S_{CaM}(t)$</td><td>$S_{CaN-dp}(t)$</td><td colspan="2">$S_{CaMKII}(t+1)$</td></tr>
<tr><td>0</td><td>0</td><td colspan="2">0</td></tr>
<tr><td>0</td><td>1</td><td colspan="2">0</td></tr>
<tr><td>1</td><td>0</td><td colspan="2">1</td></tr>
<tr><td>1</td><td>1</td><td colspan="2">0</td></tr>
<tr><td rowspan="5">Scaffold</td><td rowspan="5">$S_{Scaffold}(t+1) =$<br>NOT $S_{CaN-dp}(t)$ OR $S_{CaMKII}(t)$</td><td>$S_{CaN-dp}(t)$</td><td>$S_{CaMKII}(t)$</td><td colspan="2">$S_{Scaffold}(t+1)$</td></tr>
<tr><td>0</td><td>0</td><td colspan="2">0</td></tr>
<tr><td>0</td><td>1</td><td colspan="2">1</td></tr>
<tr><td>1</td><td>0</td><td colspan="2">0</td></tr>
<tr><td>1</td><td>1</td><td colspan="2">0</td></tr>
<tr><td rowspan="5">Endo</td><td rowspan="5">$S_{Endo}(t+1) =$<br>$S_{PKC-p}(t)$ AND NOT $S_{Scaffold}(t)$</td><td>$S_{PKC-p}(t)$</td><td>$S_{Scaffold}(t)$</td><td colspan="2">$S_{Endo}(t+1)$</td></tr>
<tr><td>0</td><td>0</td><td colspan="2">0</td></tr>
<tr><td>0</td><td>1</td><td colspan="2">0</td></tr>
<tr><td>1</td><td>0</td><td colspan="2">1</td></tr>
<tr><td>1</td><td>1</td><td colspan="2">0</td></tr>
<tr><td rowspan="3">SNAREs</td><td rowspan="3">$S_{SNAREs}(t+1) = S_{CaMKII}(t)$</td><td colspan="2">$S_{CaMKII}(t)$</td><td colspan="2">$S_{SNAREs}(t+1)$</td></tr>
<tr><td colspan="2">0</td><td colspan="2">0</td></tr>
<tr><td colspan="2">1</td><td colspan="2">1</td></tr>
</table>

**Supplementary Table 4. Neuropsychiatric disorders in WHO ICD-11 classification.**

| Category | Parent class | Diseases |
|---|---|---|
| **Diseases of the nervous system (DN)** | Choreiform disorders -Secondary Chorea | Huntington disease |
| | Movement disorders | Ataxic disorders |
| | Movement disorders-Parkinsonism | Parkinson disease |
| | Epilepsy or seizures | Genetic or presumed genetic syndromes primarily expressed as epilepsy |
| | Other disorders of the nervous system | lethal Encephalopathy |
| | Multiple sclerosis or other white matter disorders | Multiple sclerosis |
| | Motor neuron disease | Amyotrophic lateral sclerosis |
| **Mental, behavioral or neurodevelopmental disorders (MBND)** | Neurocognitive disorders | Dementia |
| | | Alzheimer Disease |
| | Disorders due to substance use | Addiction |
| | Schizophrenia or other primary psychotic disorders | Schizophrenia |
| | Mood disorders | Bipolar Disorder |
| | | Depressive disorder |
| | Disorders specifically associated with stress | Post-traumatic stress disorder |
| | Neurodevelopmental disorders | Autism Spectrum Disorder |
| | Neurodevelopmental disorders | Attention Deficit Hyperactivity Disorder |
| | Personality disorders and related traits | Personality disorder, severity unspecified |
| **Developmental anomalies** | Complete trisomies of the autosomes | Down's syndrome/ Complete trisomy 21 |
| | Syndromic genetic deafness | Fraser syndrome |
| | Sex chromosome anomalies | Fragile X chromosome |
| | Syndromes with multiple structural anomalies, without predominant body system involvement | Noonan syndrome |
| | Conditions with disorders of intellectual development as a relevant clinical feature | non-syndromic X-linked intellectual disability |

**Supplementary Table 5. Related diseases and disorders.**

| Nodes | Gene name | Diseases and disorders (from literature and Database） | |
|---|---|---|---|
| | | **Diseases (DN)** | **Disorders (MBND)** |
| CaM | CALM1 | Huntington’s Disease (M (50), -- (51, 52)) | |
| CaN | PPP3CA(B) | 1. Down's syndrome (*RCAN1, ++ (53-56))–(Developmental anomalies)<br>2. Huntington’s Disease (-- (57))<br>3. Epilepsy (-- (58)) / (ClinVar)<br>4. Parkinson Diseases (-- (59))<br>5. Multiple sclerosis (GWAS) | 1. Alzheimer Disease (-- (60-63), S (64), ++(65, 66); *RCAN1 ++ (53, 67-70))<br>2. Schizophrenia (M (71, 72), S (73); *RCAN1 M (74))<br>3. Autism Spectrum Disorders (ClinVar) |
| IP3(R) | ITPR1 | 1. Huntington's disease (-- (75-78))<br>2. Autosomal dominant cerebellar ataxia (ClinVar)<br>3. Spinocerebellar ataxias (-- (75, 76, 79-81), mKO (82))/(ClinVar)<br>4. Epilepsy (++ (83-86), mKO (82)) | 1. Alzheimer disease (++ (75, 76, 87))<br>2. Schizophrenia (GWAS)<br>3. Autism Spectrum Disorders (-- (88-90))<br>4. Bipolar Disorder (++ (91-93), S (94)) |
| PKC | PRCKA (B, G) | 1. Parkinson Diseases (++ (95-98))<br>2. Amyotrophic lateral sclerosis (GWAS)<br>3. Spinocerebellar ataxias (++ (99-102), M ((103-106)))<br>4. Epilepsy (GWAS) | 1. Neuro-developmental disorder (ClinVar)<br>2. Alzheimer disease(++ (103, 107-111),-- (112-114), M (103, 115, 116))<br>3. Dementia (-- (107), mKO (117))<br>4. Bipolar Disorder (++ (118, 119)) /(GWAS)<br>5. Alcohol Addiction (GWAS) |
| PICK1 | PICK1 | | 1. Alzheimer disease (++ (120, 121))<br>2. Alcohol Addiction (++ (122)) |
| CaMKII | CAMK2A | 1. Arthrogryposis, cleft palate, craniosynostosis, and impaired intellectual development (ClinVar)<br>2. Epilepsy (++ (58, 123, 124)) | 1. Dementia (M (125))/(ClinVar)<br>2. Schizophrenia(-- (124, 126), M (127))<br>3. Alzheimer Disease(M (128)) / (ClinVar, GWAS)<br>4. Bipolar Disorder (++ (129), -- (130))<br>5. Major Depression (++ (131), -- (124)) |

| | | | |
|---|---|---|---|
| | | | 6. Post-traumatic stress disorder (m++-KO (132))<br>7. Autism Spectrum Disorders (++ (133, 134)) /(ClinVar)<br>8. Addiction (++ (135, 136), M (137)) |
| Sap97 | DLG1 | | 1. Alzheimer Disease (-- (138))<br>2. Schizophrenia (-- (139, 140), M (141-143),S (140, 144-147)) |
| Sap102 | DLG3 | Non-syndromic X-linked intellectual disability (ClinGen) - DN | 1. Dementia (M (148, 149))/ (ClinVar)<br>2. Bipolar Disorders (-- (150-152))<br>3. Schizophrenia (-- (151))<br>4. Major Depression(-- (151)) |
| NSF | NSF | 1. Parkinson's disease (GWAS)<br>2. Developmental and epileptic encephalopathy (ClinVar) | Schizophrenia (M (59, 141, 153, 154), -- (155)) |
| PP1 | PPP1CA(B) | 1. Parkinson Diseases (M (156))<br>2. Noonan Syndrome (GWAS) | Autism Spectrum Disorders(-- (157)) |
| I-1 | PPP1R1C | | Neuro-developmental disorder (ClinVar) |
| NOS | NOS1 | | 1. Alzheimer Disease (-- (158, 159))<br>2. Schizophrenia(M (160)) |
| Thorase | ATAD1 | 1. Parkinson (-- (161))<br>2. Epilepsy (M (162)) | Dementia (M (163)) |
| TARP | CACNG2 | 1. Fragile X syndrome (GWAS) - (Developmental anomalies)<br>2. Epilepsy (M (164))/(ClinVar) | 1. Dementia (-- (123), mKO (165)) /(ClinVar)<br>2. Addiction (++ (166, 167), mKO (168))/(GWAS)<br>3. Schizophrenia (-- (169-171))<br>4. Major Depression Disorders (-- (172, 173))<br>5. Bipolar Disorders (-- (171), ++ (174), S (174-176))<br>6. Attention Deficit Hyperactivity Disorder (S (177), mKO (177))<br>7. Anti-social personality disorder (S (178)) |

| | | | |
|---|---|---|---|
| PSD95 | DLG4 | 1. Parkinson (-- (179))<br>2. Epilepsy (-- (180)) | 1. Autism Spectrum Disorders (M (181-184), ++ (133, 182, 185)) / (ClinVar)<br>2. Dementia (-- (179, 180, 184, 186), mKO (187) )<br>3. Schizophrenia (-- (126, 151, 182, 188), M (143, 182, 189-192), -- (180), S (193, 194))<br>4. Bipolar Disorders (-- (151)) |
| GRIP | GRIP1 | 1. Fraser Syndrome (GWAS) - (Developmental anomalies) | 1. Intellectual developmental disorder (ClinVar)<br>2. Autism Spectrum Disorders (S (195))<br>3. Schizophrenia (++ (196, 197))<br>4. Addiction (++ (198)) |

Note:

1. *RCAN1: Regulator of Calcineurin 1, a key protein in the calmodulin pathway that suppresses calcineurin activity.
2. Genomic and psychiatric findings are annotated using the following abbreviations: M, mutant forms; --, gene loss, reduced expression, or regulatory deficiency; ++, gene overexpression or elevated activity; S, single-nucleotide polymorphisms (SNPs); mKO, knockout experiments in mice.
3. For classification of diseases and disorders into the DN and MBND groups (right two columns), see Methods and Materials, and Table S4.
4. Parenthesized numbers indicate cited literature (see Supplementary References); databases used (e.g., GWAS Catalog, ClinVar, ClinGen) are also noted.

### Supplementary References

1. Niswender CM, Conn PJ (2010): Metabotropic glutamate receptors: Physiology, pharmacology, and disease. Annu Rev Pharmacol Toxicol. 50:295-322.
2. Hammond C (2024): *Cellular and Molecular Neurophysiology*. 5 ed.: Academic Press.
3. Mikoshiba K (2007): The IP3 receptor/Ca2+ channel and its cellular function. Biochem Soc Symp. 74:9-22.
4. Luo LQ (2020): *Principles of Neurobiology(2nd ed.)*. Garland Science.
5. Angelova PR, Abramov AY (2018): Role of mitochondrial ROS in the brain: from physiology to neurodegeneration. FEBS Lett. 592:692-702.
6. Bayer KU, Schulman H (2019): CaM kinase: still inspiring at 40. Neuron. 103:380-394.
7. Mulkey RM, Endo S, Shenolikar S, Malenka RC (1994): Involvement of a Calcineurin/inhibitor-1 phosphatase cascade in hippocampal long-term depression. Nature. 369:486-488.
8. Reyes-García SE, Escobar ML (2021): Calcineurin participation in Hebbian and homeostatic plasticity associated with extinction. Front Cell Neurosci. 15:685838.
9. Wang X, Culotta VC, Klee CB (1996): Superoxide dismutase protects calcineurin from inactivation. Nature. 383:434-437.
10. Ferri A, Gabbianelli R, Casciati A, Paolucci E, Rotilio G, Carrì MT (2000): Calcineurin activity is regulated both by redox compounds and by mutant familial amyotrophic lateral sclerosis-superoxide dismutase. J Neurochem. 75:606-613.
11. Namgaladze D, Hofer HW, Ullrich V (2002): Redox control of calcineurin bytargeting the binuclear Fe2+-Zn2+ center at the enzyme active site. J Biol Chem. 277:5962-5969.
12. Callender Julia A, Newton Alexandra C (2017): Conventional protein kinase C in the brain: 40 years later. Neuronal Signaling. 1:NS20160005.
13. Newton AC (1995): Protein kinase C: Structure, function, and regulation. J Biol Chem. 270:28495-28498.
14. Perez JL, Khatri L, Chang C, Srivastava S, Osten P, Ziff EB (2001): Pick1 targets activated Protein Kinase Cα to AMPA receptor clusters in spines of hippocampal neurons and reduces surface levels of the AMPA-type glutamate receptor subunit 2. J Physiol. 21:5417-5428.
15. Xia J, Chung HJ, Wihler C, Huganir RL, Linden DJ (2000): Cerebellar long-term depression requires PKC-regulated interactions between GluR2/3 and PDZ domain–containing proteins. Neuron. 28:499-510.
16. Osten P, Khatri L, Perez JL, Köhr G, Giese G, Daly C, et al. (2000): Mutagenesis reveals a role for ABP/GRIP binding to GluR2 in synaptic surface accumulation of the AMPA receptor. Neuron. 27:313-325.
17. Matsuda S, Mikawa S, Hirai H (1999): Phosphorylation of serine-880 in GluR2 by protein kinase C prevents its C terminus from binding with glutamate receptor-interacting protein. J Neurochem. 73:1765-1768.
18. Dong H, O'Brien RJ, Fung ET, Lanahan AA, Worley PF, Huganir RL (1997): GRIP: a synaptic PDZ domain-containing protein that interacts with AMPA receptors. Nature. 386:279-284.
19. Hanley JG, Khatri L, Hanson PI, Ziff EB (2002): NSF ATPase and α-/β-SNAPs disassemble the AMPA receptor-PICK1 complex. Neuron. 34:53-67.
20. Nishimune A, Isaac JTR, Molnar E, Noel J, Nash SR, Tagaya M, et al. (1998): NSF binding to GluR2 regulates synaptic transmission. Neuron. 21:87-97.
21. Haas A (1998): NSF-fusion and beyond. Trends Cell Biol. 8:471-473.
22. Lee SH, Liu L, Wang YT, Sheng M (2002): Clathrin adaptor AP2 and NSF interact with

overlapping sites of GluR2 and play distinct roles in AMPA receptor trafficking and hippocampal LTD. Neuron. 36:661-674.

23. Zhang J, Wang Y, Chi Z, Keuss Matthew J, Pai Y-Min E, Kang HC, et al. (2011): The AAA+ ATPase thorase regulates AMPA receptor-dependent synaptic plasticity and behavior. Cell. 145:284-299.
24. Umanah GKE, Ghasemi M, Yin X, Chang M, Kim JW, Zhang J, et al. (2020): AMPA receptor surface expression is regulated by S-nitrosylation of thorase and transnitrosylation of NSF. Cell Rep. 33:108329.
25. Förstermann U, Sessa WC (2012): Nitric oxide synthases: regulation and function. Eur Heart J. 33:829-837.
26. Barbano PE, Spivak M, Flajolet M, Nairn AC, Greengard P, Greengard L (2007): A mathematical tool for exploring the dynamics of biological networks. Proc Natl Acad Sci USA. 104:19169-19174.
27. Connor JH, Quan H, Oliver C, Shenolikar S (1998): Inhibitor-1, a regulator of protein phosphatase 1 function. In: Ludlow JW, editor. *Protein Phosphatase Protocols*. Totowa, NJ: Humana Press, pp 41-58.
28. Watanabe T, Huang H-B, Horiuchi A, da Cruze Silva EF, Hsieh-Wilson L, Allen PB, et al. (2001): Protein phosphatase 1 regulation by inhibitors and targeting subunits. Proc Natl Acad Sci USA. 98:3080-3085.
29. Yasuda R, Hayashi Y, Hell JW (2022): CaMKII: a central molecular organizer of synaptic plasticity, learning and memory. Nat Rev Neurosci. 23:666-682.
30. Shioda N, Fukunaga K (2018): Physiological and pathological roles of CaMKII-PP1 signaling in the brain. Int J Mol Sci. 19:20.
31. Leonard AS, Davare MA, Horne MC, Garner CC, Hell JW (1998): SAP97 is associated with the α-amino-3-hydroxy-5-methylisoxazole-4-propionic acid receptor GluR1 subunit. J Biol Chem. 273:19518-19524.
32. De Los Reyes DA, Karkoutly MY, Zhang Y (2023): Synapse-associated protein 102 – a highly mobile MAGUK predominate in early synaptogenesis. Front Mol Neurosci. 16:1286134.
33. Liu M, Shi R, Hwang H, Han KS, Wong MH, Ren X, et al. (2018): SAP102 regulates synaptic AMPAR function through a CNIH-2-dependent mechanism. J Neurophysiol. 120:1578-1586.
34. Rumbaugh G, Sia GM, Garner CC, Huganir RL (2003): Synapse-associated protein-97 isoform-specific regulation of surface AMPA receptors and synaptic function in cultured neurons. J Neurosci. 23:4567-4576.
35. Nakagawa T, Futai K, Lashuel HA, Lo I, Okamoto K, Walz T, et al. (2004): Quaternary structure, protein dynamics, and synaptic function of SAP97 controlled by L27 domain interactions. Neuron. 44:453-467.
36. Opazo P, Labrecque S, Tigaret CM, Frouin A, Wiseman PW, De Koninck P, et al. (2010): CaMKII triggers the diffusional trapping of surface AMPARs through phosphorylation of stargazin. Neuron. 67:239-252.
37. Chen L, Chetkovich DM, Petralia RS, Sweeney NT, Kawasaki Y, Wenthold RJ, et al. (2000): Stargazin regulates synaptic targeting of AMPA receptors by two distinct mechanisms. Nature. 408:936-943.
38. Tomita S, Sekiguchi M, Wada K, Nicoll RA, Bredt DS (2006): Stargazin controls the pharmacology of AMPA receptor potentiators. Proc Natl Acad Sci USA. 103:10064-10067.
39. Sumioka A, Yan D, Tomita S (2010): TARP phosphorylation regulates synaptic AMPA receptors through lipid bilayers. Neuron. 66:755-767.
40. Vallejo D, Inestrosa NC (2018): PSD-95 (Postsynaptic Density Protein-95). In: Choi S, editor. *Encyclopedia of Signaling Molecules*. Cham: Springer International Publishing, pp 4263-

4269.

41. Kim MJ, Futai K, Jo J, Hayashi Y, Cho K, Sheng M (2007): Synaptic accumulation of PSD-95 and synaptic function regulated by phosphorylation of serine-295 of PSD-95. Neuron. 56:488-502.

42. Doherty GJ, McMahon HT (2009): Mechanisms of endocytosis. Annu Rev Biochem. 78:857-902.

43. Fiuza M, Rostosky CM, Parkinson GT, Bygrave AM, Halemani N, Baptista M, et al. (2017): PICK1 regulates AMPA receptor endocytosis via direct interactions with AP2 α-appendage and dynamin. J Cell Biol. 216:3323-3338.

44. Tomita S, Stein V, Stocker TJ, Nicoll RA, Bredt DS (2005): Bidirectional synaptic plasticity regulated by phosphorylation of stargazin-like TARPs. Neuron. 45:269-277.

45. Passafaro M, Piëch V, Sheng M (2001): Subunit-specific temporal and spatial patterns of AMPA receptor exocytosis in hippocampal neurons. Nat Neurosci. 4:917-926.

46. Wu H, Nash JE, Zamorano P, Garner CC (2002): Interaction of SAP97 with minus-end-directed actin motor myosin VI: Implications for AMPA receptor trafficking. J Biol Chem. 277:30928-30934.

47. Jurado S (2014): The dendritic SNARE fusion machinery involved in AMPARs insertion during long-term potentiation. Front Cell Neurosci. 8:407.

48. Collingridge GL, Isaac JTR, Wang YT (2004): Receptor trafficking and synaptic plasticity. Nat Rev Neurosci. 5:952-962.

49. Kennedy MJ, Ehlers MD (2006): Organelles and trafficking machinery for postsynaptic plasticity. Annu Rev Neurosci. 29:325-362.

50. Dudek NL, Dai Y, Muma NA (2010): Neuroprotective effects of calmodulin peptide 76-121aa: disruption of calmodulin binding to mutant Huntingtin. Brain Pathol. 20:176-189.

51. Dudek NL, Dai Y, Muma NA (2008): Protective effects of interrupting the binding of calmodulin to mutant Huntingtin. J Neuropathol Exp. 67:355-365.

52. Bao J, Sharp AH, Wagster MV, Becher M, Schilling G, Ross CA, et al. (1996): Expansion of polyglutamine repeat in huntingtin leads to abnormal protein interactions involving calmodulin. Proc Natl Acad Sci USA. 93:5037-5042.

53. Harris CD, Ermak G, Davies KJA (2005): Multiple roles of the DSCR1 (Adapt78 or RCAN1) gene and its protein product Calcipressin 1 (or RCAN1) in disease. Cell Mol Life Sci. 62:2477-2486.

54. Sun X, Wu Y, Chen B, Zhang Z, Zhou W, Tong Y, et al. (2011): Regulator of calcineurin 1 (RCAN1) facilitates neuronal apoptosis through caspase-3 activation. J Biol Chem. 286:9049-9062.

55. Arron JR, Winslow MM, Polleri A, Chang C-P, Wu H, Gao X, et al. (2006): NFAT dysregulation by increased dosage of DSCR1 and DYRK1A on chromosome 21. Nature. 441:595-600.

56. Baek K-H, Zaslavsky A, Lynch RC, Britt C, Okada Y, Siarey RJ, et al. (2009): Down's syndrome suppression of tumour growth and the role of the calcineurin inhibitor DSCR1. Nature. 459:1126-1130.

57. Costa V, Giacomello M, Hudec R, Lopreiato R, Ermak G, Lim D, et al. (2010): Mitochondrial fission and cristae disruption increase the response of cell models of Huntington's disease to apoptotic stimuli. EMBO Mol Med. 2:490-503.

58. Lie AA, Blumcke L, Beck H, Schramm J, Wiestler OD, Elger CE (1998): Altered patterns of Ca2+/calmodulin-dependent protein kinase II and calcineurin immunoreactivity in the hippocampus of patients with temporal lobe epilepsy. J Neuropathol Exp Neurol. 57:1078-1088.

59. Taymans J-M, Baekelandt V (2014): Phosphatases of α-synuclein, LRRK2, and tau:

important players in the phosphorylation-dependent pathology of Parkinsonism. Front Genet. 5:382.

60. Pei JJ, Sersen E, Iqbal K, Grundkeiqbal I (1994): Expression of protein phosphatases (PP-1, PP-2A, PP-2B and PTP-1B) and protein kinases (MAP kinase and P34(CDC2)) in the hippocampus of patients with Alzheimer disease and normal aged individuals. Brain Res. 655:70-76.

61. Abdul HM, Sama MA, Furman JL, Mathis DM, Beckett TL, Weidner AM, et al. (2009): Cognitive decline in Alzheimer's disease is associated with selective changes in calcineurin/NFAT signaling. J Neurosci. 29:12957-12969.

62. Sun B, Halabisky B, Zhou Y, Palop JJ, Yu G, Mucke L, et al. (2009): Imbalance between GABAergic and glutamatergic transmission impairs adult neurogenesis in an animal model of Alzheimer's disease. Cell Stem Cell. 5:624-633.

63. Harris KA, Oyler GA, Doolittle GM, Vincent I, Lehman RAW, Kincaid RL, et al. (1993): Okadaic acid induces hyperphosphorylated forms of tau protein in human brain slices. Ann Neurol. 33:77-87.

64. Cruchaga C, Kauwe JSK, Mayo K, Spiegel N, Bertelsen S, Nowotny P, et al. (2010): SNPs associated with cerebrospinal fluid phospho-tau levels influence rate of decline in Alzheimer's disease. PLos Genet. 6:e1001101.

65. Hata R, Masumura M, Akatsu H, Li F, Fujita H, Nagai Y, et al. (2001): Up-regulation of calcineurin A beta mRNA in the Alzheimer's disease brain: Assessment by cDNA microarray. Biochem Biophys Res Commun. 284:310-316.

66. Popugaeva E, Pchitskaya E, Bezprozvanny I (2017): Dysregulation of neuronal calcium homeostasis in Alzheimer's disease - A therapeutic opportunity? Biochem Biophys Res Commun. 483:998-1004.

67. Poppek D, Keck S, Ermak G, Jung T, Stolzing A, Ullrich O, et al. (2006): Phosphorylation inhibits turnover of the tau protein by the proteasome: influence of RCAN1 and oxidative stress. Biochem J. 400:511-520.

68. Lloret A, Badia M-C, Giraldo E, Ermak G, Alonso M-D, Pallardo FV, et al. (2011): Amyloid-beta toxicity and tau hyperphosphorylation are linked via RCAN1 in Alzheimer's disease. J Alzheimers Dis. 27:701-709.

69. Ermak G, Morgan TE, Davies KJA (2001): Chronic overexpression of the calcineurin inhibitory gene DSCR1 (Adapt78) is associated with Alzheimer's disease. J Biol Chem. 276:38787-38794.

70. Blanchard JW, Bula M, Davila-Velderrain J, Akay LA, Zhu L, Frank A, et al. (2020): Reconstruction of the human blood–brain barrier in vitro reveals a pathogenic mechanism of APOE4 in pericytes. Nat Med. 26:952-963.

71. Gerber DJ, Hall D, Miyakawa T, Demars S, Gogos JA, Karayiorgou M, et al. (2003): Evidence for association of schizophrenia with genetic variation in the 8p21.3 gene, PPP3CC, encoding the calcineurin gamma subunit. Proc Natl Acad Sci USA. 100:8993-8998.

72. Görlach J, Fox DS, Cutler NS, Cox GM, Perfect JR, Heitman J (2000): Identification and characterization of a highly conserved calcineurin binding protein, CBP1/calcipressin, in Cryptococcus neoformans. The EMBO Journal. 19:3618-3629.

73. Yamada K, Gerber DJ, Iwayama Y, Ohnishi T, Ohba H, Toyota T, et al. (2007): Genetic analysis of the calcineurin pathway identifies members of the EGR gene family, specifically EGR3, as potential susceptibility candidates in schizophrenia. Proc Natl Acad Sci USA. 104:2815-2820.

74. Tam GWC, van de Lagemaat LN, Redon R, Strathdee KE, Croning MDR, Malloy MP, et al. (2010): Confirmed rare copy number variants implicate novel genes in schizophrenia. Biochem Soc Trans. 38:445-451.

75. Fedorenko OA, Popugaeva E, Enomoto M, Stathopulos PB, Ikura M, Bezprozvanny I (2014): Intracellular calcium channels: Inositol-1,4,5-trisphosphate receptors. Eur J Pharmacol. 739:39-48.

76. Egorova PA, Bezprozvanny IB (2018): Inositol 1,4,5-trisphosphate receptors and neurodegenerative disorders. The FEBS Journal. 285:3547-3565.

77. Tang T-S, Tu H, Chan EYW, Maximov A, Wang Z, Wellington CL, et al. (2003): Huntingtin and huntingtin-associated protein 1 influence neuronal calcium signaling mediated by inositol-(1,4,5) triphosphate receptor type 1. Neuron. 39:227-239.

78. Kaltenbach LS, Romero E, Becklin RR, Chettier R, Bell R, Phansalkar A, et al. (2007): Huntingtin interacting proteins are genetic modifiers of neurodegeneration. PLos Genet. 3:e82.

79. Novak MJU, Sweeney MG, Li A, Treacy C, Chandrashekar HS, Giunti P, et al. (2010): An ITPR1 gene deletion causes spinocerebellar ataxia 15/16: A genetic, clinical and radiological description. Movement Disorders. 25:2176-2182.

80. van de Leemput J, Wavrant-De Vrièze F, Rafferty I, Bras JM, Giunti P, Fisher EMC, et al. (2010): Sequencing analysis of the ITPR1 gene in a pure autosomal dominant spinocerebellar ataxia series. Movement Disorders. 25:771-773.

81. Sasaki M, Ohba C, Iai M, Hirabayashi S, Osaka H, Hiraide T, et al. (2015): Sporadic infantile-onset spinocerebellar ataxia caused by missense mutations of the inositol 1,4,5-triphosphate receptor type 1 gene. J Neurol. 262:1278-1284.

82. Matsumoto M, Nakagawa T, Inoue T, Nagata E, Tanaka K, Takano H, et al. (1996): Ataxia and epileptic seizures in mice lacking type 1 inositol 1,4,5-trisphosphate receptor. Nature. 379:168-171.

83. DeLorenzo RJ, Sun DA, Blair RE, Sombati S (2007): An in vitro model of Stroke-Induced Epilepsy: Elucidation of The roles of Glutamate and Calcium in The induction and Maintenance of Stroke-Induced Epileptogenesis. *International Review of Neurobiology*: Academic Press, pp 59-84.

84. Mirza N, Appleton R, Burn S, Carr D, Crooks D, du Plessis D, et al. (2015): Identifying the biological pathways underlying human focal epilepsy: from complexity to coherence to centrality. Hum Mol Genet. 24:4306-4316.

85. Nagarkatti N, Deshpande LS, DeLorenzo RJ (2008): Levetiracetam Inhibits both ryanodine and IP3 receptor activated calcium induced calcium release in hippocampal neurons in culture. Neurosci Lett. 436:289-293.

86. Pal S, Sun D, Limbrick D, Rafiq A, DeLorenzo RJ (2001): Epileptogenesis induces long-term alterations in intracellular calcium release and sequestration mechanisms in the hippocampal neuronal culture model of epilepsy. Cell Calcium. 30:285-296.

87. Cheung K-H, Mei L, Mak D-OD, Hayashi I, Iwatsubo T, Kang DE, et al. (2010): Gain-of-function enhancement of IP3 receptor modal gating by familial Alzheimer’s disease–linked presenilin mutants in human cells and mouse neurons. Sci Signaling. 3:ra22.

88. Gilman Sarah R, Iossifov I, Levy D, Ronemus M, Wigler M, Vitkup D (2011): Rare de novo variants associated with autism implicate a large functional network of genes involved in formation and function of synapses. Neuron. 70:898-907.

89. Ma W-J, Hashii M, Munesue T, Hayashi K, Yagi K, Yamagishi M, et al. (2013): Non-synonymous single-nucleotide variations of the human oxytocin receptor gene and autism spectrum disorders: a case–control study in a Japanese population and functional analysis. Mol Autism. 4:22.

90. Schmunk G, Boubion BJ, Smith IF, Parker I, Gargus JJ (2015): Shared functional defect in IP3R-mediated calcium signaling in diverse monogenic autism syndromes. Transl Psychiatry. 5:e643.

91. Berridge MJ (2014): Calcium signalling and psychiatric disease: bipolar disorder and schizophrenia. Cell Tissue Res. 357:477-492.

92. Machado-Vieira R, Pivovarova NB, Stanika RI, Yuan P, Wang Y, Zhou R, et al. (2011): The Bcl-2 gene polymorphism rs956572AA increases inositol 1,4,5-trisphosphate receptor–mediated endoplasmic reticulum calcium release in subjects with bipolar disorder. Biol Psychiatry. 69:344-352.

93. Mathews R, Li PP, Young LT, Kish SJ, Warsh JJ (1997): Increased Gαq/11 immunoreactivity in postmortem occipital cortex from patients with bipolar affective disorder. Biol Psychiatry. 41:649-656.

94. Distelhorst CW, Bootman MD (2011): Bcl-2 interaction with the inositol 1,4,5-trisphosphate receptor: Role in Ca2+ signaling and disease. Cell Calcium. 50:234-241.

95. Do Van B, Gouel F, Jonneaux A, Timmerman K, Gelé P, Pétrault M, et al. (2016): Ferroptosis, a newly characterized form of cell death in Parkinson's disease that is regulated by PKC. Neurobiol Dis. 94:169-178.

96. Kaleli HN, Ozer E, Kaya VO, Kutlu O (2020): Protein kinase C isozymes and autophagy during neurodegenerative disease progression. Cells. 9:553.

97. Monteleone L, Speciale A, Valenti GE, Traverso N, Ravera S, Garbarino O, et al. (2021): PKCα inhibition as a strategy to sensitize neuroblastoma stem cells to etoposide by stimulating ferroptosis. Antioxidants. 10:691.

98. Mansour HM, Mohamed AF, El-Khatib AS, Khattab MM (2023): Kinases control of regulated cell death revealing druggable targets for Parkinson's disease. Ageing Res Rev. 85:101841.

99. Wong MMK, Hoekstra SD, Vowles J, Watson LM, Fuller G, Németh AH, et al. (2018): Neurodegeneration in SCA14 is associated with increased PKCγ kinase activity, mislocalization and aggregation. Acta Neuropathol Commun. 6:99.

100. Lambert J-C, Heath S, Even G, Campion D, Sleegers K, Hiltunen M, et al. (2009): Genome-wide association study identifies variants at CLU and CR1 associated with Alzheimer's disease. Nat Genet. 41:1094-1099.

101. Schrenk K, Kapfhammer JP, Metzger F (2002): Altered dendritic development of cerebellar Purkinje cells in slice cultures from protein kinase Cγ-deficient mice. Neuroscience. 110:675-689.

102. Adachi N, Kobayashi T, Takahashi H, Kawasaki T, Shirai Y, Ueyama T, et al. (2008): Enzymological analysis of mutant protein kinase Cγ causing spinocerebellar ataxia type 14 and dysfunction in Ca2+ homeostasis. J Biol Chem. 283:19854-19863.

103. Lordén G, Newton Alexandra C (2021): Conventional protein kinase C in the brain: repurposing cancer drugs for neurodegenerative treatment? Neuronal Signaling. 5:NS20210036.

104. Pilo CA, Baffi TR, Kornev AP, Kunkel MT, Malfavon M, Chen D-H, et al. (2022): Mutations in protein kinase Cγ promote spinocerebellar ataxia type 14 by impairing kinase autoinhibition. Sci Signaling. 15:eabk1147.

105. Shirafuji T, Shimazaki H, Miyagi T, Ueyama T, Adachi N, Tanaka S, et al. (2019): Spinocerebellar ataxia type 14 caused by a nonsense mutation in the PRKCG gene. Mol Cell Neurosci. 98:46-53.

106. Pilo CA, Newton AC (2022): Two sides of the same coin: protein kinase C γ in cancer and neurodegeneration. Front Cell Dev Biol. 10:929510.

107. Sun M-K, Alkon DL (2014): Chapter two - The "memory kinases": Roles of PKC isoforms in signal processing and memory formation. In: Khan ZU, Muly EC, editors. *Prog Mol Biol Transl Sci*: Academic Press, pp 31-59.

108. Szallasi Z, Smith CB, Pettit GR, Blumberg PM (1994): Differential regulation of protein kinase

C isozymes by bryostatin 1 and phorbol 12-myristate 13-acetate in NIH 3T3 fibroblasts. J Biol Chem. 269:2118-2124.

109. Alfonso SI, Callender JA, Hooli B, Antal CE, Mullin K, Sherman MA, et al. (2016): Gain-of-function mutations in protein kinase Cα (PKCα) may promote synaptic defects in Alzheimer's disease. Sci Signaling. 9:ra47.

110. Callender JA, Yang Y, Lordén G, Stephenson NL, Jones AC, Brognard J, et al. (2018): Protein kinase Cα gain-of-function variant in Alzheimer's disease displays enhanced catalysis by a mechanism that evades down-regulation. Proc Natl Acad Sci USA. 115:E5497-E5505.

111. Morshed N, Lee MJ, Rodriguez FH, Lauffenburger DA, Mastroeni D, White FM (2021): Quantitative phosphoproteomics uncovers dysregulated kinase networks in Alzheimer's disease. Nat Aging. 1:550-565.

112. Talman V, Pascale A, Jäntti M, Amadio M, Tuominen RK (2016): Protein kinase C activation as a potential therapeutic strategy in Alzheimer's disease: is there a role for embryonic lethal abnormal vision-like proteins? Basic Clin Physiol Pharmacol. 119:149-160.

113. Wang H-Y, Pisano MR, Friedman E (1994): Attenuated protein kinase C activity and translocation in Alzheimer's Disease brain. Neurobiol Aging. 15:293-298.

114. Lucke-Wold BP, Turner RC, Logsdon AF, Simpkins JW, Alkon DL, Smith KE, et al. (2015): Common mechanisms of Alzheimer's disease and ischemic stroke: The role of protein kinase C in the progression of age-related neurodegeneration. J Alzheimers Dis. 43:711-724.

115. Prokopenko D, Morgan SL, Mullin K, Hofmann O, Chapman B, Kirchner R, et al. (2021): Whole-genome sequencing reveals new Alzheimer's disease–associated rare variants in loci related to synaptic function and neuronal development. Alzheimer's & Dementia. 17:1509-1527.

116. Masliah E, Cole G, Hansen L, Mallory M, Albright T, Terry R, et al. (1991): Protein kinase C alteration is an early biochemical marker in Alzheimer's disease. J Physiol. 11:2759-2767.

117. Wang F, Chang G, Geng X (2014): NGF and TERT co-transfected BMSCs improve the restoration of cognitive impairment in vascular dementia rats. PLoS One. 9:e98774.

118. Manji HK, Lenox RH (1999): Protein kinase C signaling in the brain: molecular transduction of mood stabilization in the treatment of manic-depressive illness. Biol Psychiatry. 46:1328-1351.

119. Chen G, Masana MI, Manji HK (2000): Lithium regulates PKC-mediated intracellular cross-talk and gene expression in the CNS in vivo. Bipolar Disord. 2:217-236.

120. Alfonso S, Kessels HW, Banos CC, Chan TR, Lin ET, Kumaravel G, et al. (2014): Synapto-depressive effects of amyloid beta require PICK1. Eur J Neurosci. 39:1225-1233.

121. Lin EYS, Silvian LF, Marcotte DJ, Banos CC, Jow F, Chan TR, et al. (2018): Potent PDZ-domain PICK1 inhibitors that modulate amyloid beta-mediated synaptic dysfunction. Sci Rep. 8:13438.

122. Thorsen TS, Madsen KL, Rebola N, Rathje M, Anggono V, Bach A, et al. (2010): Identification of a small-molecule inhibitor of the PICK1 PDZ domain that inhibits hippocampal LTP and LTD. Proc Natl Acad Sci USA. 107:413-418.

123. Chang PK, Verbich D, McKinney RA (2012): AMPA receptors as drug targets in neurological disease--advantages, caveats, and future outlook. Eur J Neurosci. 35:1908-1916.

124. Robison AJ (2014): Emerging role of CaMKII in neuropsychiatric disease. Trends Neurosci. 37:653-662.

125. Kury S, van Woerden GM, Besnard T, Onori MP, Latypova X, Towne MC, et al. (2017): De novo mutations in protein kinase genes CAMK2A and CAMK2B cause intellectual disability. Am J Hum Genet. 101:768-788.

126. Matas E, William DJF, Toro CT (2021): Abnormal expression of post-synaptic proteins in

prefrontal cortex of patients with schizophrenia. Neurosci Lett. 745:135629.

127. Yamasaki N, Maekawa M, Kobayashi K, Kajii Y, Maeda J, Soma M, et al. (2008): Alpha-CaMKII deficiency causes immature dentate gyrus, a novel candidate endophenotype of psychiatric disorders. Mol Brain. 1:6.

128. Tu S, Okamoto S-i, Lipton SA, Xu H (2014): Oligomeric Aβ-induced synaptic dysfunction in Alzheimer's disease. Mol Neurodegener. 9:48.

129. Molnar M, Potkin SG, Bunney WE, Jones EG (2003): mRNA expression patterns and distribution of white matter neurons in dorsolateral prefrontal cortex of depressed patients differ from those in schizophrenia patients. Biol Psychiatry. 53:39-47.

130. Xing GQ, Russell S, Hough C, O'Grady J, Zhang L, Yang ST, et al. (2002): Decreased prefrontal CaMKII alpha mRNA in bipolar illness. Neuroreport. 13:501-505.

131. Novak G, Seeman P, Tallerico T (2006): Increased expression of calcium/calmodulin-dependent protein kinase II beta in frontal cortex in schizophrenia and depression. Synapse. 59:61-68.

132. Hazra S, Hazra JD, Bar-On RA, Duan Y, Edut S, Cao X, et al. (2022): The role of hippocampal CaMKII in resilience to trauma-related psychopathology. Neurobiol Stress. 21:100506.

133. Kim KC, Kim P, Go HS, Choi CS, Park JH, Kim HJ, et al. (2013): Male-specific alteration in excitatory post-synaptic development and social interaction in pre-natal valproic acid exposure model of autism spectrum disorder. J Neurochem. 124:832-843.

134. Rhee J, Park K, Kim KC, Shin CY, Chung C (2018): Impaired hippocampal synaptic plasticity and enhanced excitatory transmission in a novel animal model of autism spectrum disorders with telomerase reverse transcriptase overexpression. Mol Cells. 41:486-494.

135. Oliver RJ, Brigman JL, Bolognani F, Allan AM, Neisewander JL, Perrone-Bizzozero NI (2018): Neuronal RNA-binding protein HuD regulates addiction-related gene expression and behavior. Genes Brain and Behav. 17:e12454.

136. Mueller CP, Quednow BB, Lourdusamy A, Kornhuber J, Schumann G, Giese P (2016): CaM kinases: From memories to addiction. Trends Pharmacol Sci. 37:153-166.

137. Mijakowska Z, Lukasiewicz K, Ziolkowska M, Lipinski M, Trabczynska A, Matuszek Z, et al. (2017): Autophosphorylation of alpha isoform of calcium/calmodulin-dependent kinase II regulates alcohol addiction-related behaviors. Addict Biol. 22:331-341.

138. Marcello E, Epis R, Saraceno C, Gardoni F, Borroni B, Cattabeni F, et al. (2012): SAP97-mediated local trafficking is altered in Alzheimer disease patients' hippocampus. Neurobiol Aging. 33.

139. Toyooka K, Iritani S, Makifuchi T, Shirakawa O, Kitamura N, Maeda K, et al. (2002): Selective reduction of a PDZ protein, SAP-97, in the prefrontal cortex of patients with chronic schizophrenia. J Neurochem. 83:797-806.

140. Xu X, Liang C, Lv D, Yin J, Luo X, Fu J, et al. (2018): Association of the synapse-associated protein 97 (SAP97) gene polymorphism with neurocognitive function in schizophrenic patients. Front Psychiatry. 9:458.

141. Curtis D, Consortium UK (2016): Practical experience of the application of a weighted burden test to whole exome sequence data for obesity and schizophrenia. Ann Hum Genet. 80:38-49.

142. Hammond JC, McCullumsmith RE, Haroutunian V, Meador-Woodruff JH (2011): Endosomal trafficking of AMPA receptors in frontal cortex of elderly patients with schizophrenia. Schizophr Res. 130:260-265.

143. Xing J, Kimura H, Wang C, Ishizuka K, Kushima I, Arioka Y, et al. (2016): Resequencing and association analysis of six PSD-95-related genes as possible susceptibility genes for schizophrenia and autism spectrum disorders. Sci Rep. 6:27491.

144. Uezato A, Yamamoto N, Jitoku D, Haramo E, Hiraaki E, Iwayama Y, et al. (2017): Genetic and molecular risk factors within the newly identified primate-specific exon of the SAP97/DLG1 gene in the 3q29 schizophrenia-associated locus. Am J Med Genet B. 174:798-807.

145. Sato J, Shimazu D, Yamamoto N, Nishikawa T (2008): An association analysis of synapse-associated protein 97 (SAP97) gene in schizophrenia. J Neural Transm. 115:1355-1365.

146. Soler J, Fananas L, Parellada M, Krebs M-O, Rouleau GA, Fatjo-Vilas M (2018): Genetic variability in scaffolding proteins and risk for schizophrenia and autism-spectrum disorders: a systematic review. J Psychiatr Neurosci. 43:223-244.

147. Xu X, Wang Y, Zhou X, Yin J, Yu H, Wen X, et al. (2020): The genetic variations in SAP97 gene and the risk of schizophrenia in the Chinese Han population: a further study. Psychiatr Genet. 30:110-118.

148. Zanni G, van Esch H, Bensalem A, Saillour Y, Poirier K, Castelnau L, et al. (2010): A novel mutation in the DLG3 gene encoding the synapse-associated protein 102 (SAP102) causes non-syndromic mental retardation. Neurogenetics. 11:251-255.

149. Tarpey P, Parnau J, Blow M, Woffendin H, Bignell G, Cox C, et al. (2004): Mutations in the DLG3 gene cause nonsyndromic X-linked mental retardation. Am J Hum Genet. 75:318-324.

150. Beneyto M, Meador-Woodruff JH (2008): Lamina-specific abnormalities of NMDA receptor-associated postsynaptic protein transcripts in the prefrontal cortex in schizophrenia and bipolar disorder. Neuropsychopharmacol. 33:2175-2186.

151. Clinton SM, Meador-Woodruff JH (2004): Abnormalities of the NMDA receptor and associated intracellular molecules in the thalamus in schizophrenia and bipolar disorder. Neuropsychopharmacol. 29:1353-1362.

152. McCullumsmith RE, Kristiansen LV, Beneyto M, Scarr E, Dean B, Meador-Woodruff JH (2007): Decreased NR1, NR2A, and SAP102 transcript expression in the hippocampus in bipolar disorder. Brain Res. 1127:108-118.

153. Vine AE, McQuillin A, Bass NJ, Pereira A, Kandaswamy R, Robinson M, et al. (2009): No evidence for excess runs of homozygosity in bipolar disorder. Psychiatr Genet. 19:165-170.

154. Lencz T, Lambert C, DeRosse P, Burdick KE, Morgan TV, Kane JM, et al. (2007): Runs of homozygosity reveal highly penetrant recessive loci in schizophrenia. Proc Natl Acad Sci USA. 104:19942-19947.

155. Siegert S, Seo J, Kwon EJ, Rudenko A, Cho S, Wang W, et al. (2015): The schizophrenia risk gene product miR-137 alters presynaptic plasticity. Nat Neurosci. 18:1008-1016.

156. Edler MC, Salek AB, Watkins DS, Kaur H, Morris CW, Yamamoto BK, et al. (2018): Mechanisms regulating the association of protein phosphatase 1 with spinophilin and neurabin. ACS Chem Neurosci. 9:2701-2712.

157. Li J, Chai A, Wang L, Ma Y, Wu Z, Yu H, et al. (2015): Synaptic P-Rex1 signaling regulates hippocampal long-term depression and autism-like social behavior. Proc Natl Acad Sci USA. 112:E6964-E6972.

158. Bonnar O, Hall CN (2020): First, tau causes NO problem. Nat Neurosci. 23:1035-1036.

159. Aissa BM, Lee HS, Bennett MB, Thatcher RJG (2016): Targeting NO/cGMP signaling in the CNS for neurodegeneration and Alzheimer’s disease. Curr Med Chem. 23:2770-2788.

160. Candemir E, Kollert L, Weissflog L, Geis M, Mueller A, Post AM, et al. (2016): Interaction of NOS1AP with the NOS-I PDZ domain: Implications for schizophrenia-related alterations in dendritic morphology. Eur Neuropsychopharmacol. 26:741-755.

161. Gao F, Zhang H, Yang J, Cai M, Yang Q, Wang H, et al. (2022): ATPase Thorase deficiency causes alpha-synucleinopathy and Parkinson's disease-like behavior. Cells. 11:2990.

162. Ahrens-Nicklas RC, Umanah GKE, Sondheimer N, Deardorff MA, Wilkens AB, Conlin LK, et

al. (2017): Precision therapy for a new disorder of AMPA receptor recycling due to mutations in ATAD1. Neurol Genet. 3:e130.

163. Umanah GKE, Pignatelli M, Yin X, Chen R, Crawford J, Neifert S, et al. (2017): Thorase variants are associated with defects in glutamatergic neurotransmission that can be rescued by Perampanel. Sci Transl Med. 9:eaah4985.

164. Weston MC (2017): Two targets are better than one: A new strategy to increase the specificity of anti-epileptic drugs. Epilepsy Curr. 17:235-236.

165. Caldeira GL, Inacio AS, Beltrao N, Barreto CAV, Rodrigues MV, Rondao T, et al. (2022): Aberrant hippocampal transmission and behavior in mice with a stargazin mutation linked to intellectual disability. Mol Psychiatry. 27:2457-2469.

166. Hoffman JL, Faccidomo S, Saunders BL, Taylor SM, Kim M, Hodge CW (2021): Inhibition of AMPA receptors (AMPARs) containing transmembrane AMPAR regulatory protein γ-8 with JNJ-55511118 shows preclinical efficacy in reducing chronic repetitive alcohol self-administration. Alcohol Clin Exp Res. 45:1424-1435.

167. Kato AS, Witkin JM (2018): Protein complexes as psychiatric and neurological drug targets. Biochem Pharmacol. 151:263-281.

168. Faccidomo SP, Homan JL, Agoglia AE, Taylor SM, Kim M, Herman MA, et al. (2021): Transmembrane AMPA regulatory protein-gamma 8 (TARP) modulation of AMPA receptor regulation of alcohol reinforcement and excitatory transmission in the BLA. Alcohol Clin Exp Res. 45:87A.

169. Drummond JB, Tucholski J, Haroutunian V, Meador-Woodruff JH (2013): Transmembrane AMPA receptor regulatory protein (TARP) dysregulation in anterior cingulate cortex in schizophrenia. Schizophr Res. 147:32-38.

170. Benesh JL, Mueller TM, Meador-Woodruff JH (2022): AMPA receptor subunit localization in schizophrenia anterior cingulate cortex. Schizophr Res. 249:16-24.

171. Gill MB, Bredt DS (2011): An emerging role for TARPs in neuropsychiatric disorders. Neuropsychopharmacol. 36:362-363.

172. Woo HI, Lim S-W, Myung W, Kim DK, Lee S-Y (2018): Differentially expressed genes related to major depressive disorder and antidepressant response: genome-wide gene expression analysis. Exp Mol Med. 50:1–11.

173. Liu H, Wang P, Song W, Sun X (2009): Degradation of regulator of calcineurin 1 (RCAN1) is mediated by both chaperone-mediated autophagy and ubiquitin proteasome pathways. FASEB J. 23:3383-3392.

174. Silberberg G, Levit A, Collier D, Clair DS, Munro J, Kerwin RW, et al. (2008): Stargazin involvement with bipolar disorder and response to lithium treatment. Pharmacogenet Genomics. 18:403-412.

175. Nissen S, Liang S, Shehktman T, Kelsoe JR, Bipolar Genome Study Bi GS (2012): Evidence for association of bipolar disorder to haplotypes in the 22q12.3 region near the genes stargazin, IFT27 and parvalbumin. Am J Med Genet B. 159B:941-950.

176. Miranda A, Shekhtman T, McCarthy M, DeModena A, Leckband SG, Kelsoe JR (2019): Study of 45 candidate genes suggests CACNG2 may be associated with lithium response in bipolar disorder. J Affect Disord. 248:175-179.

177. Bai W-J, Luo X-G, Jin B-H, Zhu K-S, Guo W-Y, Zhu X-Q, et al. (2022): Deficiency of transmembrane AMPA receptor regulatory protein gamma-8 leads to attention-deficit hyperactivity disorder-like behavior in mice. Zool Res. 43:851-870.

178. Peng S-X, Wang Y-Y, Zhang M, Zang Y-Y, Wu D, Pei J, et al. (2021): SNP rs10420324 in the AMPA receptor auxiliary subunit TARP γ-8 regulates the susceptibility to antisocial personality disorder. Sci Rep. 11:11997.

179. Whitfield DR, Vallortigara J, Alghamdi A, Howlett D, Hortobágyi T, Johnson M, et al. (2014): Assessment of ZnT3 and PSD95 protein levels in Lewy body dementias and Alzheimer's disease: association with cognitive impairment. Neurobiol Aging. 35:2836-2844.

180. Fernandez E, Collins MO, Frank RAW, Zhu F, Kopanitsa MV, Nithianantharajah J, et al. (2017): Arc requires PSD95 for assembly into postsynaptic complexes involved with neural dysfunction and intelligence. Cell Rep. 21:679-691.

181. Quintero-Rivera F, Sharifi-Hannauer P, Martinez-Agosto JA (2010): Autistic and psychiatric findings associated with the 3q29 microdeletion syndrome: Case report and review. Am J Med Genet Part A. 152A:2459-2467.

182. Coley AA, Gao W-J (2018): PSD95: A synaptic protein implicated in schizophrenia or autism? Prog Neuro-Psychopharmacol Biol Psychiatry. 82:187-194.

183. Santini E, Klann E (2014): Reciprocal signaling between translational control pathways and synaptic proteins in autism spectrum disorders. Sci Signaling. 7:re10.

184. Nithianantharajah J, Komiyama NH, McKechanie A, Johnstone M, Blackwood DH, Clair DS, et al. (2013): Synaptic scaffold evolution generated components of vertebrate cognitive complexity. Nat Neurosci. 16:16-24.

185. Schütt J, Falley K, Richter D, Kreienkamp HJ, Kindler S (2009): Fragile X mental retardation protein regulates the levels of scaffold proteins and glutamate receptors in postsynaptic densities. J Biol Chem. 284:25479-25487.

186. Chen H, Qiao D, Wang C, Zhang B, Wang Z, Tang L, et al. (2022): Fragile X mental retardation protein mediates the effects of androgen on hippocampal PSD95 expression and dendritic spines density/morphology and autism-like behaviors through miR-125a. Front Cell Neurosci. 16:872347.

187. Wang W, Zhang X-y, Feng Z-g, Wang D-x, Zhang H, Sui B, et al. (2017): Overexpression of phosphodiesterase-4 subtypes involved in surgery-induced neuroinflammation and cognitive dysfunction in mice. Brain Res Bull. 130:274-282.

188. MacDonald ML, Banerjee A, Ciccimaro E, Blair IA, Hahn C-G (2011): Altered intracellular trafficking of PSD proteins in postmortem brains of schizophrenia. Biol Psychiatry. 69:13S.

189. Funk AJ, Mielnik CA, Koene R, Newburn E, Ramsey AJ, Lipska BK, et al. (2017): Postsynaptic density-95 isoform abnormalities in schizophrenia. Schizophr Bull. 43:891-899.

190. Matosin N, Fernandez-Enright F, Lum JS, Engel M, Andrews JL, Gassen NC, et al. (2016): Molecular evidence of synaptic pathology in the CA1 region in schizophrenia. npj Schizophr. 2:16022.

191. Brennand KJ, Simone A, Jou J, Gelboin-Burkhart C, Tran N, Sangar S, et al. (2011): Modelling schizophrenia using human induced pluripotent stem cells. Nature. 473:221-225.

192. Soler J, Fañanás L, Parellada M, Krebs M-O, Rouleau GA, Fatjó-Vilas M (2018): Genetic variability in scaffolding proteins and risk for schizophrenia and autism-spectrum disorders: a systematic review. J Psychiatry Neurosci. 43:223-244.

193. Li J-M, Lu C-L, Cheng M-C, Luu S-U, Hsu S-H, Chen C-H (2013): Exonic resequencing of the DLGAP3 gene as a candidate gene for schizophrenia. Psychiatry Research. 208:84-87.

194. Cheng M-C, Lu C-L, Luu S-U, Tsai H-M, Hsu S-H, Chen T-T, et al. (2010): Genetic and functional analysis of the DLG4 gene encoding the post-synaptic density protein 95 in schizophrenia. PLoS One. 5:e15107.

195. Mejias R, Adamczyk A, Anggono V, Niranjan T, Thomas GM, Sharma K, et al. (2011): Gain-of-function glutamate receptor interacting protein 1 variants alter GluA2 recycling and surface distribution in patients with autism. Proc Natl Acad Sci USA. 108:4920-4925.

196. Hammond JC, McCullumsmith RE, Funk AJ, Haroutunian V, Meador-Woodruff JH (2010): Evidence for abnormal forward trafficking of AMPA receptors in frontal cortex of elderly

patients with schizophrenia. Neuropsychopharmacol. 35:2110-2119.

197. Dracheva S, McGurk SR, Haroutunian V (2005): mRNA expression of AMPA receptors and AMPA receptor binding proteins in the cerebral cortex of elderly schizophrenics. J Neurosci Res. 79:868-878.

198. Chen H, Chen L, Yuan Z, Yuan J, Li Y, Xu Y, et al. (2022): Glutamate receptor-interacting protein 1 in D1-and D2-dopamine receptor-expressing medium spiny neurons differentially regulates cocaine acquisition, reinstatement, and associated spine plasticity. Front Cell Neurosci. 16:979078.
